# FtsW protein-protein interactions visualized in live *Staphylococcus aureus* cells by FLIM-FRET

**DOI:** 10.1101/2025.05.30.657058

**Authors:** Nils Y. Meiresonne, Sara F. Costa, Simon Schäper, Mário J. Ferreira, Patricia Reed, Zach Hensel, Fábio Fernandes, Mariana G. Pinho

**Affiliations:** Bacterial Cell Biology, Instituto de Tecnologia Química e Biológica António Xavier, Universidade NOVA de Lisboa, Oeiras, Portugal; Single Molecule Microbiology, Instituto de Tecnologia Química e Biológica António Xavier, Universidade NOVA de Lisboa, Oeiras, Portugal; iBB-Institute for Bioengineering and Biosciences and i4HB-Institute for Health and Bioeconomy, Instituto Superior Técnico, Universidade de Lisboa, Lisboa, Portugal; Bioengineering Department, Instituto Superior Técnico, Universidade de Lisboa, Av. Rovisco Pais, Lisboa, Portugal

## Abstract

The bacterial cell cycle relies on the coordinated and dynamic interactions between division proteins and those involved in peptidoglycan (PG) synthesis. However, visualizing these interactions *in vivo* remains technically challenging. Here, we established fluorescence-lifetime imaging microscopy combined with Förster resonance energy transfer (FLIM-FRET) as a robust, spatially resolved technique to visualize protein interactions in living *Staphylococcus aureus* using fluorescent proteins. After systematically optimizing growth conditions and the analysis pipeline, we validated the method with cytosolic and membrane-anchored control proteins, achieving FRET efficiencies of up to 40%. Using FLIM-FRET, we mapped the protein interactions of the critical glycosyltransferase FtsW within the septal PG synthesizing complex. We confirmed its interaction with the cognate transpeptidase PBP1 and the regulatory protein DivIB. Notably, we found that FtsW also self-interacts, an observation corroborated by an alternative FLIM-FRET method employing Halo-Tag labelled with Janelia Fluor dyes. These findings support the hypothesis that septal PG synthesis may be carried out by a complex of multimers, capable of simultaneously synthesizing more than one glycan strand.

Inhibition of PG synthesis by directly targeting PBP1 with the beta-lactam antibiotic imipenem weakened the interaction between PBP1 and FtsW, whereas the FtsW self-interaction was enhanced in a dose-dependent manner. In contrast, inhibition of PG synthesis by targeting the lipid II flippase, therefore depleting the FtsW-PBP1 substrate from the outer surface of the cell membrane, had little effect on these interactions. This suggests that alterations in FtsW interactions result primarily from antibiotic-induced conformational changes or from uncoupling the activities of FtsW and PBP1, resulting in the presence of uncrosslinked glycans, rather than merely from a loss of PG synthesis activity.

## Introduction

Bacterial growth requires tight coordination of cell division and peptidoglycan (PG) synthesis to ensure a balance between robustness and flexibility over the cell cycle. This balance is orchestrated by a multiprotein complex called the divisome, which contains cell division proteins and PG synthesis enzymes and assembles the division septum [1– 3]. Although the composition of the divisome has been studied for some years, its mode of assembly and its dynamics are only now becoming apparent. Central to bacterial cell division, the divisome represents a promising but underexplored target for antibiotic development, particularly in pathogens like *Staphylococcus aureus*.

In *S. aureus*, as in most bacteria, division starts with midcell recruitment of the orchestrator protein FtsZ, which is tethered to the membrane through FtsA [4,5]. Here, FtsZ forms a discontinuous, dynamic ring of tubulin-like filaments that treadmill around the division plane, recruiting several early division proteins among which are the FtsZ regulators EzrA and SepF [6,7]. Subsequently, late divisome proteins DivIB, DivIC and FtsL, and PG synthesis enzymes FtsW and PBP1 are recruited and positioned to synthesize the new septum [8–11]. FtsW is a shape, elongation, division, and sporulation (SEDS) protein with glycosyltransferase activity, while PBP1 is a monofunctional penicillin binding protein with transpeptidase activity [12]. Together, they synthesize septal PG by incorporating incoming PG precursor lipid II molecules into the existing PG, through the synthesis of the *N*-Acetylglucosamine - *N*-Acetylmuramic acid (GlcNAc-MurNAc) glycan backbone by FtsW, and the crosslinking of different glycan strands via pentapeptide crossbridges by PBP1 [12,13]. FtsW consists of ten transmembrane segments and thus has an extensive surface for potential protein interactions within the membrane or on either side of the membrane bilayer. Structurally, *S. aureus* FtsW is predicted to form a complex with PBP1, DivIB, DivIC and FtsL to drive constriction of the division septum [14].

The divisome is a highly dynamic complex of proteins that function and interact with each other throughout the bacterial cell cycle. Yet, relatively little is known about the dynamics of protein-protein interactions in the divisome, some of which may be transient rather than stable. Seminal studies using fluorescence microscopy to investigate divisome formation revealed that in *Escherichia coli, Bacillus subtilis* and *S. aureus* divisome proteins assemble in a sequential manner and in two distinct steps [8,10,11]. The so-called early divisome prepares the future division site and distributes late divisome proteins that build the new septum and their regulator proteins. In fact, after an initial collaborative period, division and PG synthesis complexes appear to function independently as physically separated complexes [10,15–17]. This prompts the field to revise existing models to represent the transient character of these protein complexes. Biochemical assays alone, although important, does not fully disclose where, or at what point during the cell cycle, interactions of the divisome occur.

A powerful technique for detecting protein-protein interaction dynamics in single living cells, with spatiotemporal resolution, is Fluorescence Lifetime Imaging Microscopy-Förster Resonance Energy Transfer (FLIM-FRET) [18]. FRET occurs when compatible donor and acceptor fluorophores are in close proximity, typically within 10 nm. This is similar to the distance at which protein interactions take place and therefore measuring FRET is a valid method to study these interactions. FLIM is a reliable technique to detect FRET because fluorescence lifetime is a spectroscopic parameter independent of protein expression levels, molecular diffusion or excitation fluctuations [19]. Fluorescence lifetime is the average time an excited fluorophore stays in the excited state before returning to the ground state. If donor and acceptor fluorophores are in close enough proximity for FRET to occur, an additional relaxation pathway is provided for the donor, resulting in a shorter fluorescence lifetime. Importantly, FLIM-FRET provides information not only on whether protein interactions occur, but also on where and when they take place.

Despite its advantages, FLIM-FRET is not routinely used to probe protein interactions in bacterial cells due to its technical complexity. To date, only a few studies have applied FLIM-FRET to probe protein interactions in live bacterial cells [12,20–26]. This paper describes the implementation of FLIM-FRET as a robust technique to assess protein-protein interactions in living bacteria. Using this technique in *S. aureus* cells, we studied the interactions of proteins that are part of the septal PG synthesis complex FtsW, PBP1and DivB. Our findings show *in vivo* interactions between FtsW-PBP1, FtsW-DivIB and, interestingly, demonstrate that FtsW interacts with itself. This supports the hypothesis that multiple glycan strands are synthesized simultaneously, in a coordinated manner, by a complex with more than one FtsW molecule. Furthermore, we provide compelling evidence that the interactions of the septal PG synthesis complex undergo changes in conformation or transiency in the presence of different antibiotics. Together, our results establish FLIM-FRET as a powerful tool for assessing bacterial protein interaction states, with spatial resolution, paving the way for *in vivo* characterization of their stable or transient interactions.

## Results

### Establishing sfTq2^ox^-mNG as a FLIM-FRET pair for *S. aureus*

To implement FLIM-FRET for the study of interactions with spatiotemporal information in live *S. aureus* cells, we started by testing several experimental parameters. First, superfolder mTurquoise2ox (sfTq2^ox^) and mNeonGreen (mNG) proteins were chosen as donor/acceptor FRET pair for their favourable spectroscopic properties over other FRET pairs, and the relatively long fluorescence lifetime (τ) of ∼4 ns and monoexponential fluorescence decay of the sfTq2^ox^ donor [23,27] (**Supplementary Text 1 and Figure S1**). Second, different growth media were tested to ensure a low fluorescence background in FLIM-FRET experiments while maintaining robust growth of *S. aureus*. This resulted in the selection of EZ Rich Defined Medium supplemented with 5 % Tryptic Soy Broth (TSB) (which we will refer to as EZ-rich) over both TSB and M9 minimal medium (**Supplementary text 2 and Figure S2**). Third, an analysis pipeline was established to mathematically remove unwanted signal from potential light scattering at the glass interface of microscopy slides (**Supplementary text 3 and Figure S3)**.

Once experimental parameters were established, a set of strains was constructed as controls for validation of FLIM-FRET in the *S. aureus* strain JE2. These included (i) BCBNM002 constitutively producing sfTq2^ox^ (donor), (ii) BCBNM003 producing a tandem of mNG fused to sfTq2^ox^ through a short amino acid linker (positive control), and (iii) BCBNM005 producing unlinked sfTq2^ox^ and mNG (negative control) (**Figure 1a,b**). In all cases, genes encoding fluorescent protein (FP) were constitutively expressed from the chromosomal *spa* locus through the unrepressed xylose promoter [28].

**Figure 1.**
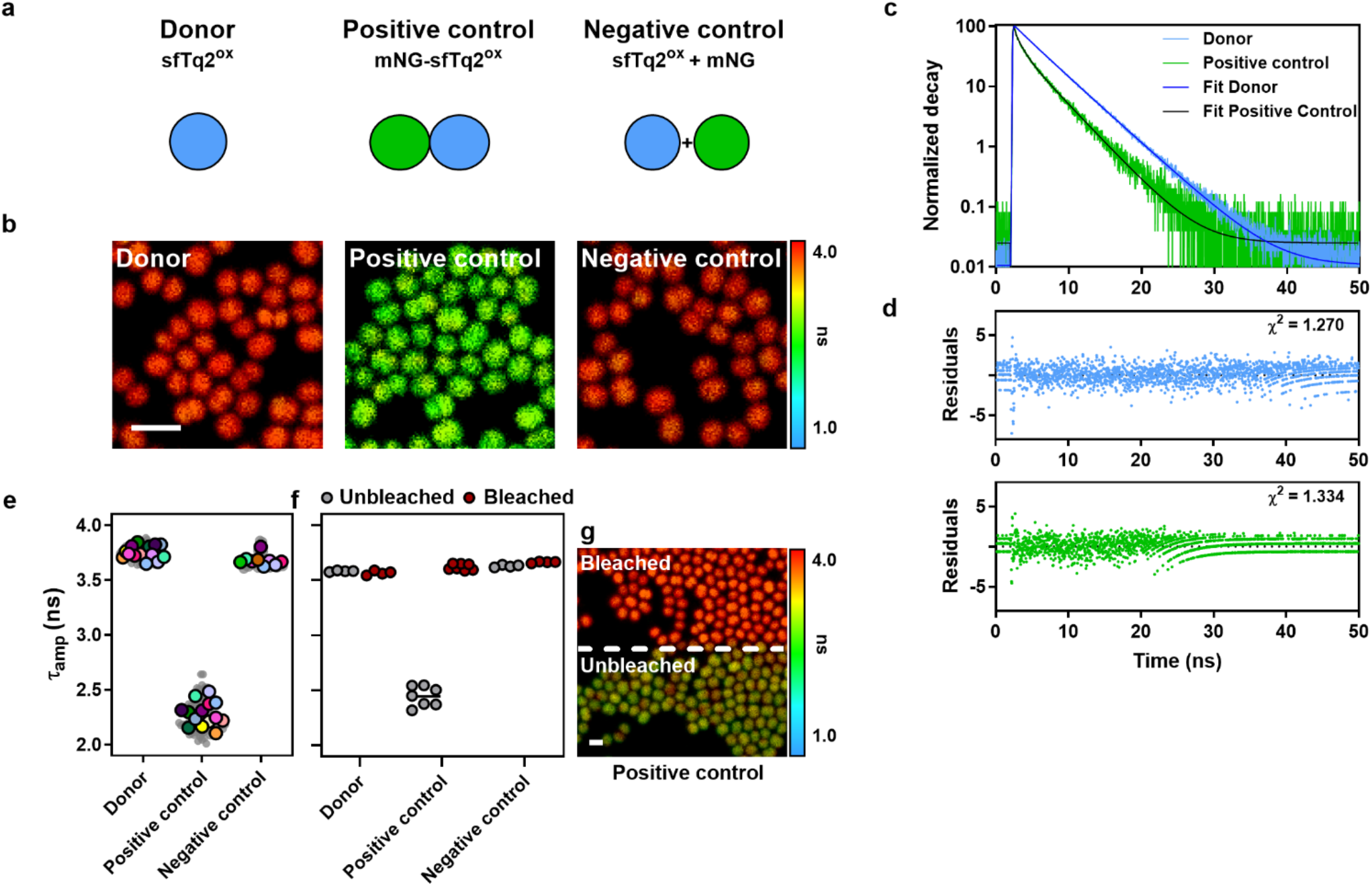
Validation of FLIM-FRET in *S. aureus* cells using cytosolic sfTq2^ox^ and mNG proteins. **a**) Cartoon of donor and acceptor fluorescent proteins used in control strains BCBNM002 (Donor), BCBNM003 (Positive control) and BCBNM005 (Negative control). **b**) FastFLIM (**see Methods**) images showing a reduction in the sfTq2^ox^ lifetime in the positive, but not in the negative control strain, when compared with the donor strain. **c**) Normalized fluorescence decay curves of sfTq2^ox^ (donor BCBNM002, blue) or sfTq2^ox^-mNG (positive control BCBNM003, green) produced in JE2 cells. **d**) Residuals of the fitted decay curves shown in c, positive control = blue, Negative control = green. Representative decay curves and fitting data are presented in **Figure S5. e**) Superplots of the amplitude weighted fluorescence lifetimes measured for the indicated stains. Each grey dot represents the fitted lifetime of a single FLIM image. Each coloured dot represents the average lifetime across FLIM images within an individual experiment. **f**) Fluorescence lifetimes before and after acceptor photobleaching for the donor, positive control and negative control strains. Comparing these values respectively indicates -0.4 %, 32.5 % and 1.0 % FRET using formula FRET efficiency = (1 − (τ_amp_ unbleached ROI /τ_amp_ Bleached ROI)) * 100. **g**) Representative FastFLIM image of the positive control strain in which the top region was bleached. All scale bars represent 2 µm.

The strains were grown to OD_600_ ∼0.4-0.5, the cells were washed three times with phosphate buffered saline (PBS), placed on microscopy slides with a 1.2 % agarose in PBS pad and fluorescence decays were measured using a confocal microscope. The fluorescence decay curves (**Figure 1c,d**) were fitted to an amplitude weighted average fluorescence lifetime (τ_amp_) weighing each lifetime component (τ_1_, τ_2_, etc.) by its amplitude to accommodate for heterogeneity and interactions with the local environment in biological systems [29]. All decay curves were fitted by a three-component reconvolution fitting model, after which the scatter component was subtracted (**Supplementary text 3, Materials and Methods**). For simplicity, from here onwards, we will refer to the τ_amp_ value obtained from fitting the fluorescence decay curves as “lifetime” of a fluorophore. The sfTq2^ox^ lifetimes measured from the control strains were unaffected by potential bleed-through of mNG fluorescence into the donor channel (**Supplementary text 4 and Figure S4**).

We then tested the ability to detect protein interactions by FLIM-FRET using the constructed control strains (**Figure 1e**). The sfTq2^ox^ lifetime in the donor strain BCBNM002 was 3.76 ± 0.06 ns (mean ± SD). As expected, sfTq2^ox^’s lifetime in the presence of unlinked mNG acceptor, obtained from negative control strain BCBNM005, was very similar (3.68 ± 0.05 ns) given that no FRET was anticipated. In contrast, lifetime of sfTq2^ox^ fused to mNG in the positive control strain BCBNM003 was substantially lower (2.28 ± 0.11 ns) and the measured fluorescence decay curve showed the characteristic decrease associated with high FRET (**Figure 1c,d**). Representative examples of fluorescence decay curves and fitting data are shown in **Figure S5**. It should be noted that the fluorescence lifetimes of sfTq2^ox^ in positive control strains, where it is physically linked to mNG, showed more variation than in strains in which no interaction occurred. This is likely due to high FRET levels reducing the number of photons detected for the sfTq2^ox^ donor, thereby decreasing the signal-to-noise-ratio. Indeed, although fluorescence lifetime is an intensity-independent parameter, its measurement still depends on the number of photons collected. FRET efficiency can be calculated from the decrease in lifetime of a sample relative to that of the reference (donor) using the formula FRET efficiency = (1 − (τ_amp_ sample/τ_amp_ reference)) * 100. Applying this formula revealed an impressive average FRET efficiency of ∼40 % for the positive control, whereas the negative control only showed ∼2 %.

An effective way to verify whether the decrease of fluorescence lifetime of a donor protein is indeed due to FRET between interacting proteins is to perform an internal control by photobleaching the acceptor protein, impairing its ability to accept energy. Upon acceptor photobleaching, the donor fluorophore loses its additional (FRET) relaxation pathway and its average (donor) lifetime is restored (*i*.*e*., increases). To demonstrate the effectiveness of the acceptor photobleaching set up, we determined that (i) the mNG acceptor can be readily bleached, (ii) without adversely affecting the sfTq2^ox^ donor, and that (iii) bleaching restores the lifetime of sfTq2^ox^ in the positive control strain to donor only levels (**Figure S6**). Specifically, a full field of view (FOV) was imaged, but the fluorescence decay curves of the bleached and unbleached regions of interest (ROIs) were analysed separately. The resulting lifetimes for the bleached and unbleached ROIs of the sfTq2^ox^ donor strain (BCBNM002) and the negative control strain (BCBNM005) were similar and close to the value obtained from the full unbleached FOV (**Figure 1f,g**). This indicates that, as expected, bleaching the mNG acceptor in these strains did not affect the sfTq2^ox^ lifetime. Strikingly, for the positive control strain producing the mNG-sfTq2^ox^ tandem (BCBNM003), the unbleached and bleached ROIs yielded respective lifetimes of 2.5 ns and 3.6 ns, indicating a recovery of the donor lifetime to values similar to the negative control strain where no FRET occurs (**Figure 1f,g)**.

The FLIM-FRET results were further validated using the independent, intensity-based method of spectral unmixing, where FRET efficiency is calculated based on the additional intensity of the acceptor fluorophore when in the presence of a donor fluorophore (sensitized emission, E*f*A) [30,31]. Fluorescence emission spectra of strains producing sfTq2^ox^ (BCBNM002), mNG (BCBNM001) or no fluorescent protein (JE2) served as references for donor-, acceptor- or background-fluorescence, respectively. These were used to unmix the fluorescence spectra of the above mentioned negative (BCBNM005) and positive (BCBNM003) control strains into their individual spectral components. This resulted in respective E*f*A values of ∼1 % and 30 %, corroborating the results obtained with these control strains in the FLIM-FRET experiments (**Figure S7**).

### Protein interactions detected by FLIM-FRET in *S. aureus*

A tandem construct in which donor and acceptor molecules are physically linked (**Figure 1a**) served as a positive control for FLIM-FRET experiments. However, it is not a realistic representation of cellular protein interactions, which occur between two independently produced proteins. To better mimic protein interactions in bacterial cells, we used the interacting Zip domains (Zip^18^ and Zip^25^) of the CyaA adenylate cyclase from *Bordetella pertussis* that are routinely used as a positive control in bacterial two-hybrid interaction assays [32,33]. For this, we genetically fused the Zip domains to the sfTq2^ox^ donor and mNG acceptor, so that the Zip-Zip interaction would bring the two fluorophores into close proximity, resulting in FRET. A set of strains was constructed that produce sfTq2^ox^-Zip^25^ (BCBNM016, donor), mNG-Zip^18^ + sfTq2^ox^-Zip^25^ (BCBNM019, positive control) or sfTq2^ox^-Zip^25^ + mNG (BCBNM020, negative control) in the *S. aureus* strain JE2, constitutively expressing the corresponding genes from the chromosomal *spa* locus through the unrepressed xylose promoter (**Figure 2a,b**)[28]. The strains were cultured as described above and the cells were harvested for FLIM. The fluorescence lifetime of sfTq2^ox^ in the donor and in the negative control strain was similar (3.69 ± 0.02 ns and 3.64 ± 0.02 ns, respectively) (**Figure 2c**). The difference in lifetimes calculated to an energy transfer efficiency of 1.4 % suggesting that, as expected, no interactions occurred, apart from potential negligible random interactions due to molecular crowding. In contrast, a shorter fluorescence lifetime of 3.19 ± 0.02 ns was measured in the positive control strain, confirming the Zip^18^-Zip^25^ interaction with an energy transfer efficiency of 13.6 %. Representative fluorescence decays curves and fitting data are presented in **Figure S8**. These results were validated by acceptor photobleaching (**Figure 2d,e**). Additional validation for the Zip^18^-Zip^25^ interaction was obtained by spectral unmixing with an E*f*A value of 16.2 ± 2.0 %, for the positive control strain (**Figure S9**). These results confirmed the negligible (or absent) interaction between donor and acceptor proteins in the negative control strain and the Zip-Zip interaction in the positive control strain.

**Figure 2.**
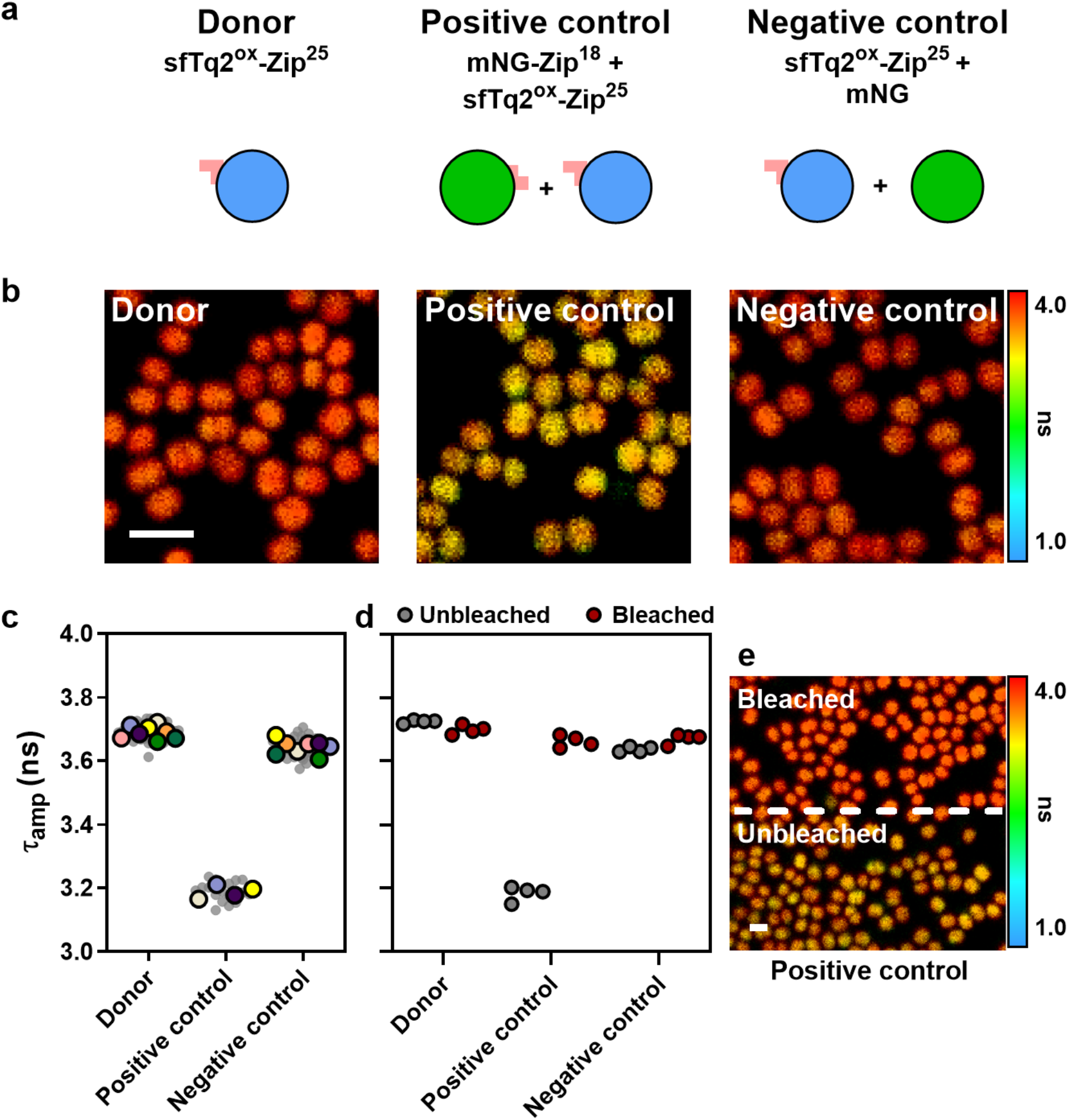
Assessing the Zip-Zip interaction in *S. aureus* by FLIM-FRET. **a**) Cartoon of Zip^18^ and Zip^25^ donor and acceptor fluorescent protein fusions produced in strains BCBNM016 (Donor), BCBNM019 (Positive control) and BCBNM020 (Negative control). **b**) FastFLIM (see Methods) images showing fluorescence lifetime differences between the donor, positive and negative control strains. **c**) Superplots of the amplitude weighted average fluorescence lifetimes measured for the indicated strains. Each grey dot represents the fitted lifetime of a single FLIM image. Each coloured dot represents the average lifetime across FLIM images within an individual experiment. Representative decay curves and fitting data are presented in **Figure S8. d**) Fluorescence lifetimes before and after acceptor photobleaching for the donor, positive control and negative control strains, respectively indicating -0.7 %, 13.1 % and 0.9 % FRET based on the difference in lifetime. **e**) Representative FastFLIM image of the positive control strain in which the top region is bleached, showing a lifetime difference in comparison to the unbleached bottom region. All scale bars represent 2 µm.

### Membrane localized protein interactions detected by FLIM-FRET in *S. aureus*

After using FLIM-FRET to visualize cytosolic protein interactions in *S. aureus*, we wanted to show that this technique can also be used to study interactions of proteins with a specific subcellular localization in live bacterial cells. For this, we fused sfTq2^ox^ and mNG-sfTq2^ox^ to the transmembrane domain (TM) of *S. aureus* PBP2 targeting the fluorescent proteins to the membrane, resulting in strains expressing the genes to produce sfTq2^ox^-TM (BCBNM010, donor) and the tandem mNG-sfTq2^ox^-TM (BCBNM011, positive control) from the *S. aureus* strain JE2 *spa* locus, under the control of the aTc-inducible *xyl-tetO* promoter (**Figure 3a**). Two negative control strains consisted of membrane localized sfTq2^ox^-TM combined with either cytosolic mNG (BCBNM015, negative control 1) or membrane localized mNG through a fusion with the full-length, 6 transmembrane domain, membrane localized protein SAUSA300_2100, referred to as Lrp, produced from its native genomic locus (BCBNM053, negative control 2). The strains were cultured and production of the fusion proteins was induced as described above. The fluorescence lifetime of sfTq2^ox^-TM when produced alone was 3.58 ± 0.04 ns. The shorter lifetime for sfTq2^ox^-TM compared to cytosolic sfTq2^ox^ and sfTq2^ox^-Zip^18^ may be explained by membrane crowding conditions, as previously observed for membrane localized fusions in *E. coli* [21]. The sfTq2^ox^ lifetime in the positive control strain producing the tandem mNG-sfTq2^ox^-TM (BCBNM011) was 2.24 ± 0.12 ns, which corresponds to 37.3 % FRET. The fluorescence lifetime of membrane sfTq2^ox^-TM localized donor was unaffected by the presence of either cytosolic or membrane localized mNG acceptor (3.60 ± 0.07 ns and 3.53 ± 0.08 ns for negative controls 1 and 2, respectively), suggesting no random interactions due to membrane crowding (**Figure 3c**). Representative fluorescence decay curves for these data are presented in **Figure S10**.

**Figure 3.**
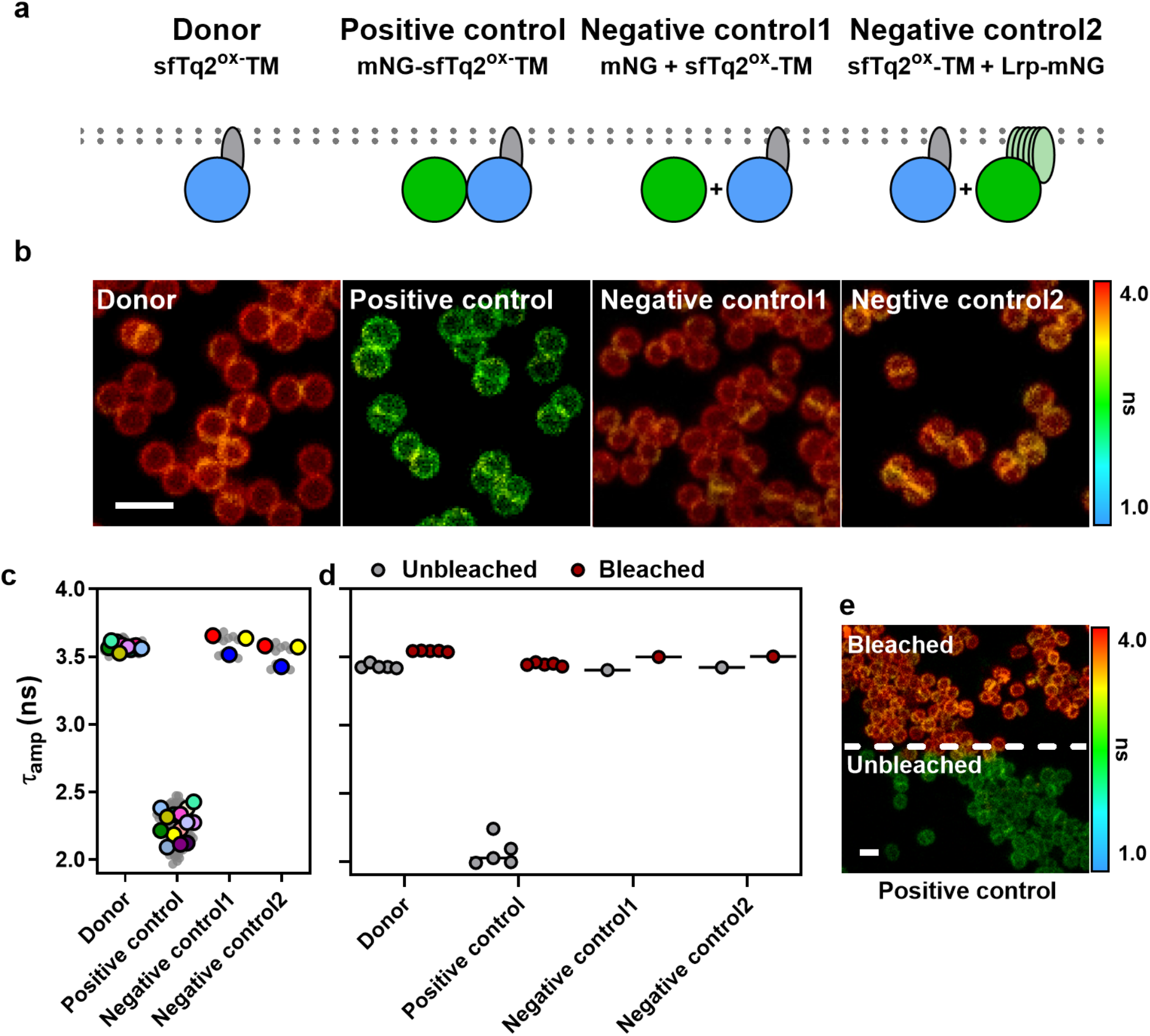
Membrane localized protein-protein interactions assessed by FLIM-FRET. **a**) Cartoon representation of donor and acceptor fluorescent protein fusions to transmembrane domains produced by strains BCBNM010 (Donor), BCBNM011 (Positive control), BCBNM015 (Negative control 1) and BCBNM053 (Negative control 2). **b**) FastFLIM images of strains described in a. **c**) Superplots of the fitted lifetimes from the FLIM measurements. Each grey dot represents the fitted lifetime of a single FLIM image. Each coloured dot represents the average lifetime across FLIM images within an individual experiment. **d**) Fluorescence lifetimes before and after acceptor photobleaching for the indicated strains, showing lifetime recovery for the acceptor photobleached positive control. **e**) Representative FastFLIM image of the positive control strain in which the top region was bleached, showing a lifetime difference in comparison to the unbleached bottom region. All scale bars represent 2 µm.

The results obtained for the membrane associated control strains were validated by acceptor photobleaching, which restored the fluorescence lifetime of sfTq2^ox^ in the positive control to 3.45 ns (**Figure 3d,e**). Further confirmation of membrane localized FRET was obtained by unmixing of the positive control strain fluorescence spectra, which resulted in E*f*A values of 30.0 ± 2.8 % for the positive control (**Figure S11**). Taken together, these results suggest the potential to detect a broad FLIM-FRET range for membrane localized protein interactions and pave the way for assessing *in vivo* protein interactions in *S. aureus* with spatiotemporal information.

### *S. aureus* FtsW protein-protein interactions assessed by FLIM-FRET

After demonstrating that FLIM-FRET can reliably detect protein interactions at the *S. aureus* membrane in the control strains, we proceeded to investigate interactions between staphylococcal proteins. We selected proteins crucial for cell cycle progression as candidates, because the ability to detect their interactions *in vivo* provides a valuable tool for functional studies and antibiotic development.

The conserved protein FtsW plays a central role in cell division, functioning in septal PG synthesis, together with its cognate partner, the class B penicillin-binding protein PBP1. Alphafold modelling predicted an interaction between *S. aureus* FtsW and PBP1, which we recently confirmed [12–14,34]. This makes the FtsW-PBP1 interaction ideal to test the implemented FLIM-FRET approach using the sfTq2^ox^-mNG FRET-pair. To investigate this interaction, we constructed a set of strains producing the protein fusions from the chromosomal *spa* region under the control of the aTc-inducible *xyl-tetO* promoter. Central to these experiments was donor strain BCBNM023 which produces FtsW-sfTq2^ox^ and served as a reference for strains co-expressing mNG-tagged potential interaction partners. The fluorescence lifetime of FtsW-sfTq2^ox^ in the donor strain was 3.55 ± 0.07 ns. Coproduction of FtsW-sfTq2^ox^ with mNG-PBP1 in strain BCBNM031 resulted in a decrease of sfTq2^ox^ lifetime to 3.00 ± 0.06 ns, corresponding to a FLIM-FRET efficiency of 15.5 %, confirming, *in vivo*, the interaction between FtsW and PBP1 (**Figure 4a,b**).

**Figure 4.**
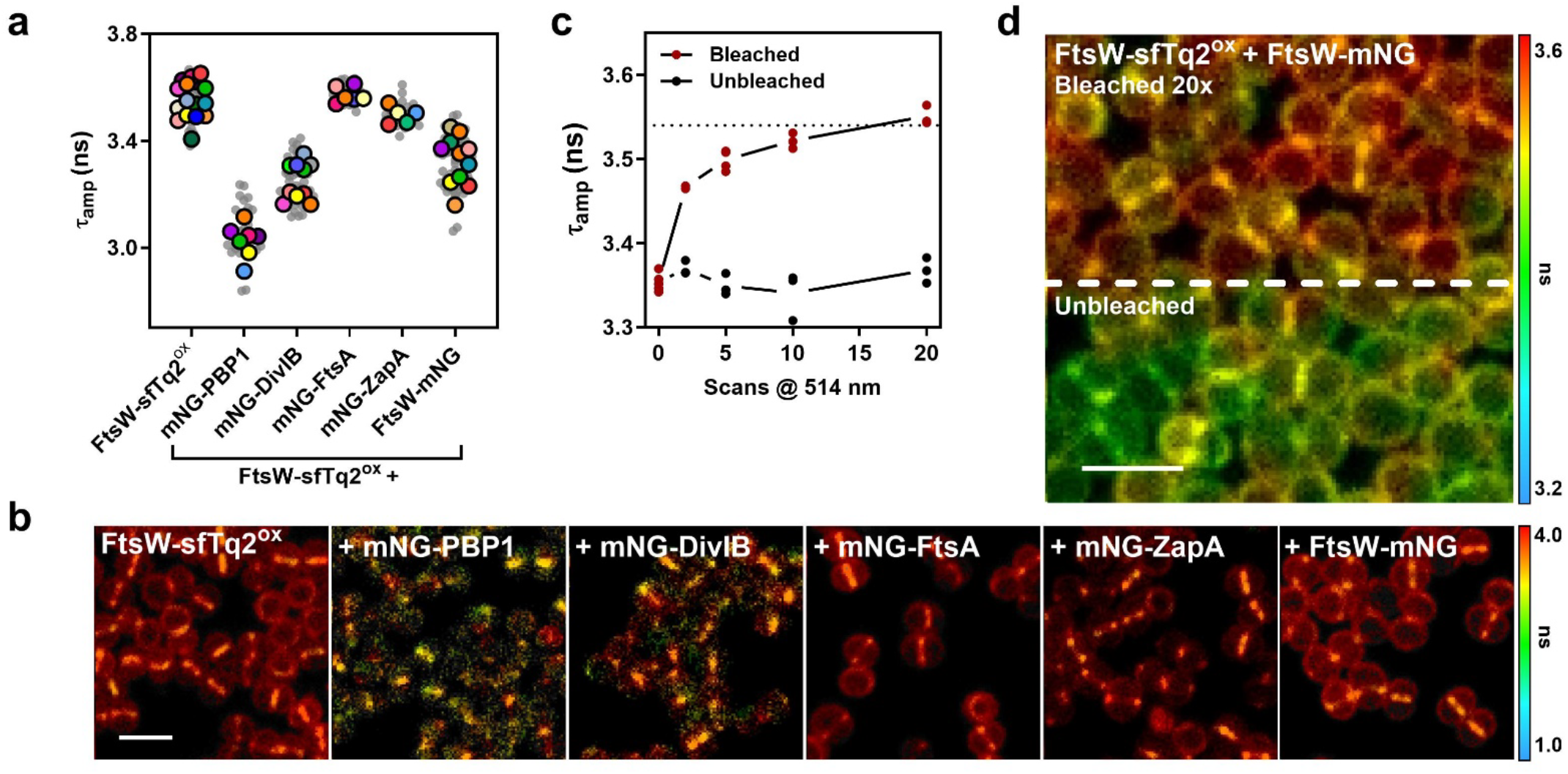
FtsW protein-protein interactions with divisome proteins measured by FLIM-FRET. **a**) Superplots of the fluorescence lifetimes measured for the indicated stains producing FtsW-sfTq2^ox^ (BCBNM023), + mNG-PBP1 (BCBNM031), + mNG-DivIB (BCBNM037), +mNG-FtsA (BCBNM032), + mNG-ZapA (BCBNM035) and + FtsW-mNG (BCBNM033). Each grey dot represents the fitted lifetime obtained from a single FLIM image, while each coloured dot represents the average lifetime across FLIM images within an individual experiment. **b**) Representative FastFLIM (**see Methods**) images of the strains indicated in a. **c**) The effect of acceptor photobleaching in strain BCBNM033 shows the restoration of the FtsW-sfTq2^ox^ lifetime whereas the lifetime measured in the unbleached regions remains stably low. **d**) FastFLIM image of strain BCBNM033 of which the top region was acceptor photobleached by 20 scans with the 514 nm laser. The bleached (top) and unbleached (bottom) regions of the image show a clear difference in fluorescence lifetime. All scale bars represent 2µm.

FtsW-PBP1 activity must be tightly regulated to balance PG synthesis during the cell cycle, but little is known about the direct and/or transient interactions that govern this activity. Both proteins, as well as their putative activator protein DivIB, move around the division site at the same velocity [14]. Inhibition of PG synthesis with antibiotics stops the directional movement of the three proteins, suggesting direct or indirect interactions [14]. To assess the direct interaction between FtsW and DivIB, mNG-DivIB was co-produced alongside FtsW-sfTq2^ox^ (BCBNM037) resulting in a fluorescence lifetime of 3.26 ± 0.09 ns. This 0.29 ns decrease compared to the FtsW-sfTq2^ox^ lifetime of the donor strain indicates a close proximity between FtsW and DivIB for FRET to occur with 8.3 % efficiency, indicating that the two proteins likely interact, which to the best of our knowledge, has not yet been demonstrated *in vivo* **(Figure 4a,b)**.

Another putative activator of septal PG synthesis is the actin-like, membrane associated FtsZ anchor FtsA. In *E. coli*, this activation is mediated by FtsN (a protein absent in *S. aureus*), which acts through FtsA and FtsQLB (homologs of *S. aureus* DivIB-DivIC-FtsL), switching the subcomplex to an “ON” state and initiating FtsW-PBP3 activity (PBP3 is the cognate bPBP of FtsW in *E. coli*, equivalent to PBP1 in *S. aureus*) [35,36]. Although *S. aureus* lacks FtsN, bacterial two-hybrid data suggest staphylococcal FtsA and FtsW interact [37] making this pair an interesting candidate for FLIM-FRET analysis. When mNG-FtsA was coproduced with FtsW-sfTq2^ox^ in strain BCBNM032 the fluorescence lifetime was 3.57 ± 0.03 ns, nearly identical to that in the donor strain (3.55 ± 0.07 ns), suggesting no detectable interaction between FtsW and FtsA **(Figure 4a,b)**. However, it should be noted that the absence of FRET alone does not demonstrate the absence of an interaction.

ZapA, another division protein that interacts with FtsZ, but is not expected to interact with FtsW, was also assessed. Strain BCBNM035 producing mNG-ZapA and FtsW-sfTq2^ox^ exhibited a fluorescence lifetime of 3.50 ± 0.04 ns suggesting no interaction between FtsW and ZapA, as expected **(Figure 4a,b)**.

Even though the septal PG synthesis complex is often modelled as a collection of monomers, several of its protein components are known to form higher order structures. For instance, various PBPs are predicted or have been shown to function as homo-dimers, including *S. aureus* PBP1 [21,38,39]. Additionally, studies on the *E. coli* FtsQBL subcomplex suggest a potential 2:2:2 stoichiometry, forming a dimer of trimers [40–42]. In this context, the observed interactions of FtsW with PBP1 and DivIB raise the question whether FtsW also self-interacts and, more broadly, whether the divisome PG synthesis complex functions as a multimer. To assess FtsW-FtsW interaction, FtsW-mNG was coproduced with FtsW-sfTq2^ox^ in *S. aureus* (BCBNM033), resulting in a FtsW-sfTq2^ox^ fluorescence lifetime of 3.33 ± 0.09 ns. Compared to the donor strain, the observed decrease corresponds to a FRET efficiency of 6.2 %, supporting the hypothesis of FtsW self-interaction. In fact, this is most likely an underestimation of the FtsW self-interaction, as FtsW-sfTq2^ox^ donor molecules in this strain have the potential to interact with (i) unlabelled FtsW (produced from the native locus), (ii) another donor labelled FtsW or (iii) acceptor labelled FtsW, with only the latter interaction resulting in FRET. Representative fluorescence decay curves and fitting data of the strains used to assay FtsW interactions are presented in **Figure S12**.

Further support for FtsW self-interaction came from acceptor photobleaching in strain BCBNM033 producing both FtsW-sfTq2^ox^ and FtsW-mNG, which resulted in the full restoration of the FtsW-sfTq2^ox^ lifetime to values observed when the donor fluorophore is in the absence of an acceptor. Moreover, the lifetime recovery scaled with the number of bleaching cycles (**Figure 4c,d**). Additional evidence came from spectral unmixing of the fluorescence spectra measured from the strain BCBNM033 resulting in an E*f*A value of 7.1 ± 0.9 %, again strongly suggesting FRET and the presence of an FtsW-FtsW interaction (**Figure S13a,b**).

An interesting alternative approach to investigate self-interaction of proteins by FLIM-FRET was employed to assess FtsW-FtsW multimerization using Halo-tag, a self-labelling protein tag that covalently binds fluorescent ligands such as Janelia Fluor rhodamine dyes [43,44]. Compared to fluorescent proteins, these dyes have superior photophysical properties and were already used for single-molecule localization microscopy of FtsW in *S. aureus* [14]. FtsW fused Halo tag (FtsW-HT) was inducibly produced from the *spa* locus in strain BCBSS187. Labelling this strain with a 1:1 mix of donor and acceptor dyes should result, on average, in half of the FtsW-HT molecules being labelled with the donor dye and half with the acceptor dye. In these conditions each FtsW-HT protein labelled with the donor dye could partner with (i) unlabelled native FtsW, (ii) donor-labelled FtsW-HT or (iii) acceptor-labelled FtsW-HT with only the latter option able to result in FRET. Dyes JF503 (donor) and JF585 (acceptor) form a suitable FRET pair with a Förster radius of 5.8 nm and were selected for the experiments. FtsW-HT produced in strain BCBSS187 was labelled with either donor JF503 dye or with a 1:1 mix of JF503:JF585 dyes and fluorescence decay curves were measured. The JF503 donor labelled sample fitted to an average fluorescence lifetime of 3.1 ± 0.0 ns, whereas the sample labelled with the 1:1 mix of dyes exhibited a reduced lifetime of 2.7 ± 0.1 ns indicating an energy transfer rate of 12.8 % (**Figure S13c-f**). This result provides additional evidence for the FtsW self-interaction and, importantly, is a proof-of-concept for the alternative approach of using Halo-tag and JF dyes for protein self-interaction FLIM-FRET studies.

### Impact of antibiotic treatment on the FtsW-PBP1-DivIB complex

The processive movement of *S. aureus* FtsW, DivIB and PBP1 during septal PG synthesis slows down in the presence of antibiotics targeting PG synthesis [14]. We hypothesized that the interactions between these three proteins, observed by FLIM-FRET, could also be impacted by PG synthesis inhibitors, possibly revealing conformational changes.

To investigate whether antibiotic treatment affects the FtsW-PBP1 interaction, the FtsW-sfTq2^ox^ donor strain (BCBNM023) and the FtsW-sfTq2^ox^and mNG-PBP1 producing strain (BCBNM031) were treated with 1 µg/ml imipenem, a β-lactam that preferentially inhibits PBP1’s transpeptidase activity [45]. Untreated control cultures were grown in parallel for direct comparison. As expected, the fluorescence lifetime of the FtsW-sfTq2^ox^ donor strain did not change in the absence or presence of imipenem (3.6 ns in both cases) (**Figure 5a and S14**). Imipenem treatment did, however, affect the FtsW-PBP1 interaction, leading to an increase in FtsW-sfTq2^ox^ lifetime from 2.9 ns in the absence of imipenem to 3.2 ns in its presence (**Figure 5a**). This increase suggests that inhibition of PBP1 TPase domain/activity either increased the distance between the donor and acceptor fluorophores, possibly due to conformational changes, or increased transience of the interaction. In any case, this result provides evidence for activity-related changes of the FtsW-PBP1 PG synthesis complex, revealed by FLIM-FRET. We notice, however, that the fluorescence lifetime of FtsW-sfTq2^ox^ in the imipenem-treated cells was still lower than in donor strain BCBNM023, indicating that FtsW-PBP1 interaction was not completely abolished in the presence of the antibiotic.

**Figure 5.**
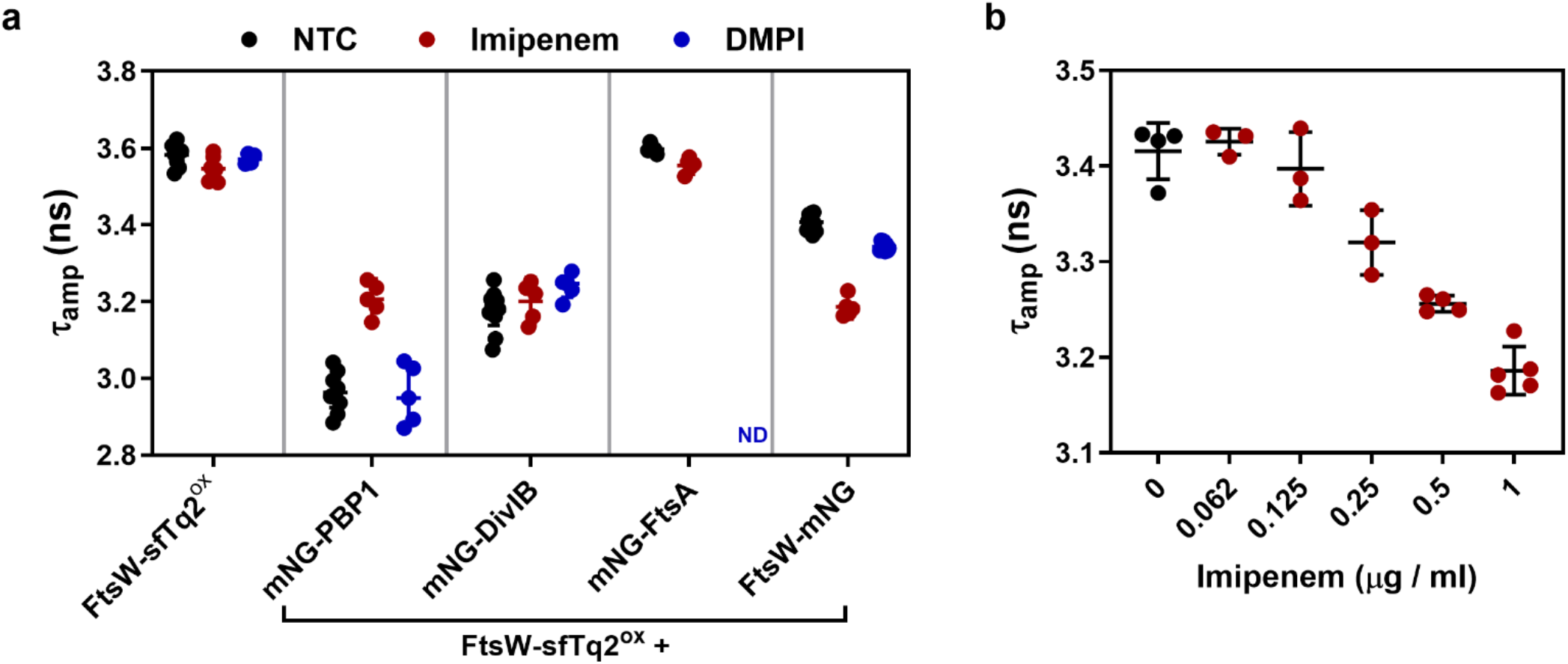
Effect of antibiotic treatment on FtsW protein-protein interactions. **a**) Fluorescence lifetimes of sfTq2^ox^ in the strains producing FtsW-sfTq2^ox^ alone (BCBNM023) or combined with mNG-PBP1 (BCBNM031), mNG-DivIB (BCBNM037), mNG-FtsA (BCBNM032), and FtsW-mNG (BCBNM033). The strains were grown without antibiotic (non-treated control, NTC, black), or in the presence of imipenem (1 µg/ml, red) or DMPI (8 µg/ml, blue). ND = not determined. **b**) The fluorescence lifetime of the FtsW-sfTq2^ox^ in the presence of FtsW-mNG (BCBNM033) decreases under increasing concentrations of imipenem. Representative fluorescence decay curves for the strains mentioned for panel a treated with imipenem or DMPI are presented in **Figure S14-16**.

Considering the higher order organization of the PG synthesizing complex, the response of the FtsW-FtsW interaction to imipenem treatment was also assessed. Strain BCBNM033 (used to assess the FtsW-FtsW interaction) was grown and exposed to a range of imipenem concentrations (0.0625 – 1 µg/ml, plus 0 µg/ml as a control). Strikingly, the fluorescence lifetimes of FtsW-sfTq2^ox^ inversely correlated with the concentration of imipenem, decreasing from ∼3.4 ns (in the absence of antibiotic) to below 3.2 ns (in the presence 1 µg/ml of imipenem). This corresponds to an increase in FRET efficiency as the antibiotic concentration increases (**Figure 5b and S15**). These results show that inhibition of PBP1 TPase activity by imipenem directly affects FtsW self-interaction or its interaction dynamics. The donor and acceptor FPs may come into closer proximity due to changes in FtsW-FtsW conformation or interaction dynamics. One hypothesis would be that blocking PBP1’s TPase crosslinking activity may alter FtsW’s ability/activity to incorporate additional glycan units, resulting in a conformation in which the protein termini are positioned closer together.

We also assessed the effects of imipenem treatment on the FtsW-DivIB interaction. Strain BCBNM037, producing FtsW-sfTq2^ox^ and mNG-DivIB, was treated with imipenem (1 µg/ml), which caused no substantial changes in the donor fluorescence lifetime compared to the non-treated control (3.25 ns and 3.20 ns, respectively) (**Figure 5a**). This suggests that the FtsW-DivIB interaction is largely independent of PBP1 activity, perhaps indicating that this interaction is decoupled from FtsW-PBP1 activity or interaction dynamics. To investigate whether inhibition of PBP1 could promote an interaction between FtsW and FtsA, strain BCBNM032 producing FtsW-sfTq2^ox^ and FtsA-mNG was treated with imipenem. As expected, no substantial changes in fluorescence lifetime were observed compared to the non-treated control (3.56 and 3.60 ns, respectively (**Figure 5a**).

The effect of imipenem on FtsW interactions could result either from the inhibition of PG synthesis or from conformational changes caused directly by the binding of the antibiotic to the transpeptidase active site of PBP1. To determine if the inhibition of PG synthesis alone was sufficient to alter FtsW interactions, we used DMPI, an antibiotic that blocks PG synthesis by inhibiting MurJ, the enzyme responsible for flipping the PG synthesis precursor lipid II across the membrane. Unlike β-lactams, DMPI does not bind PG synthases but instead depletes their substrate on the outer surface of the membrane [46,47]. Importantly, DMPI slows down the movement of FtsW and DivIB around the division septum, similar to β-lactams [14]. In contrast to imipenem, DMPI treatment did not affect the FtsW-PBP1 or FtsW-FtsW interaction when assessed by FLIM FRET (**Figure 5a and S16**). Additionally, the FtsW-DivIB interaction remained largely unaffected, showing only minor variations. These findings indicate that the observed changes in FtsW interactions caused by imipenem were not simply due to the inhibition of PG synthesis itself but instead appeared to require direct antibiotic binding to PBP1 or specific inhibition of PBP1 alone. Therefore, the observed changes FtsW interactions likely resulted from conformational shifts or from uncoupling FtsW and PBP1 activities, leading to the synthesis of uncrosslinked glycans, rather than merely reflecting a general loss of PG synthesis activity.

## Discussion

Fluorescence lifetime is a spectroscopic property of fluorophores that is strongly dependent on their local environment. Therefore, it is an excellent indicator for assessing local conditions, such as the interactions between compatible fusion proteins via FRET. However, FLIM remains an underused technique in molecular microbiology due to the requirement for specialized equipment, technical complexity and the need for stringent experimental consistency. Furthermore, achieving subcellular resolution with FLIM-FRET is challenging due to the small size of bacterial cells and the diffraction limit.

In this work, we systematically optimized a FLIM-FRET assay (**Supplementary text 1-4**) to reliably detect bacterial protein interactions. We demonstrated that FLIM-FRET is a robust tool to investigate bacterial protein interactions, both for proteins freely diffusing in the cytosol (**Figures 1 and 2**) or localized to the membrane (**Figure 3**).

An important advantage of FLIM-FRET experiments is the possibility of performing an intrinsic control by photobleaching the acceptor fluorophore within an interacting pair. This prevents energy transfer from donor to acceptor restoring the donor fluorescence lifetime to values similar to those obtained under non-interacting conditions. Comparing the lifetimes obtained from acceptor photobleached and unbleached regions within a FLIM image of a strain producing interacting proteins enables the detection of FRET using a single sample. Therefore, a well-established sensor strain is sufficient to assess an interaction of interest under different experimental conditions. However, there are limitations, as insufficient bleaching will fail to restore the donor-only lifetime, whereas excessive bleaching may introduce phototoxic effects, such as fluorophore degradation or generation of reactive oxygen species.

Once robust FLIM-FRET conditions were established, we applied this technique to investigate protein interactions of interest that are challenging to study *in vivo*. Specifically, we focused on bacterial cell division proteins, which are recognized as promising antibiotic targets for developing urgently needed novel antibiotics. Characterizing their (possibly transient) interactions is essential for understanding the dynamics of the division machinery. FtsW plays a pivotal role in bacterial division as it is the primary glycosyltransferase responsible for polymerizing the sugar-chain backbone of the septal cell wall PG, which is essential for *S. aureus* viability. FtsW directly interacts with its cognate partner, the transpeptidase PBP1 [12]. Single-molecule tracking has revealed that FtsW and PBP1 move around the division plane, together with the putative activator protein DivIB, and subsequent modelling suggests an interaction of PBP1-FtsW with the DivIB-DivIC-FtsL subcomplex [14]. Using FLIM-FRET, we confirmed the direct interaction of FtsW with PBP1 (∼16 % energy transfer) and revealed the potential interaction of FtsW with DivIB (∼8 % energy transfer) (**Figure 4**). In contrast, no interactions were observed between FtsW and cell division proteins FtsA or ZapA. However, it is important to note that while detected FRET strongly supports a direct interaction, the absence of FRET does not necessarily indicate a lack of interaction. Factors such as a large physical separation between the domains fused to the donor and acceptor fluorophore or unfavourable dipole orientation could prevent FRET detection despite an actual protein interaction.

An interesting finding was the FtsW self-interaction. Although the associated energy transfer was not very high (∼6 %), this result is reliable, as it was validated by (i) acceptor photobleaching, (ii) spectral unmixing and (iii) FLIM-FRET using a FtsW-HaloTag fusion labelled with the JF503-JF585 dye FRET pair (∼13 % energy transfer) (**Figure S13**). This result indicates that the PG synthesis complex is not necessarily a molecular machine of strictly monomers, but rather that its individual components are present as self-interacting multimers, resulting in the synthesis and crosslinking of more than one glycan strand at a time. This hypothesis is supported by the fact that PBPs and the DivIB-DivIC-FtsL subcomplex have been shown to self-interact in *S. aureus* and other bacteria [21,23,37–42]. It is also in line with the hypothetical multienzyme complex proposed by J-V. Höltje over 25 years ago to explain how *E. coli* could grow its stress-bearing PG sacculus by a safe mechanism, inserting three new glycan strands for each strand that is removed [48,49]. This “three-for-one” mechanism implies that three glycan strands would be synthesized simultaneously.

Interestingly, inhibition of PBP1’s active site with imipenem destabilized the FtsW-PBP1 complex (**Figure 5**). This initially suggested that an active PBP1 might be required for the FtsW-PBP1 interaction to achieve maximum FRET efficiency, potentially due to a favourable protein conformation or decreased transience of the interaction. However, this hypothesis appears unlikely, as inhibiting PG synthesis by blocking the translocation of its lipid II with DMPI—a condition under which PBP1 lacks its substrate—did not significantly affect the fluorescence lifetime of the donor protein used to measure the FtsW-PBP1 interaction. The key difference between these two inhibitory conditions is that, in the presence of imipenem, PBP1 is directly targeted by the antibiotic, possibly inducing conformational changes. Additionally, in these conditions, FtsW may continue to synthesize glycan strands, but PBP1 is unable to crosslink them. This scenario may lead to the accumulation of long, uncrosslinked glycans in the space between the membrane and the existing PG layer, potentially causing deleterious effects and destabilizing the FtsW-PBP1 complex. In contrast, DMPI treatment prevents substrate availability on the outer side of the membrane, thus inhibiting glycan synthesis by FtsW altogether (and consequently also preventing crosslinking by PBP1). Under these conditions, despite impaired PG synthesis, the FtsW-PBP1 interaction and the FtsW self-interaction remained unaffected. These mechanistic details can only be assessed in live cells, showing the power of FLIM-FRET to detect dynamic changes in protein interactions as a result of changes in the environment.

## Material and methods

### Strains and growth conditions

All strains used in this study are listed in **Table S1**. *E. coli* strains DC10B or DH5α were used for cloning and plasmid propagation and were grown in LB (Miller) medium (VWR) supplemented with 100 µg / ml Ampicillin (Sigma) at 37 °C. *S. aureus* strains were grown in Tryptic Soy Broth (TSB, VWR) at 37 °C or 30 °C for all experiments except FLIM assays. For FLIM experiments, cells were grown in MOPS EZ Rich Defined Medium (Teknova Inc, M2105), supplemented with 5 % TSB, which we refer to as EZ-rich.

### Plasmid construction

All plasmids and cloning strategies, and primers used in this study are listed in **Tables S2 and S3**. Plasmids pET28a-mTq2 and pET28a-mTq2^ox^, containing *S. aureus* codon optimized mTq2 and sfTq2^ox^, were ordered from NZYtech custom gene synthesis service. The DNA templates for further cloning of *sftq2ox* and *mNG* were, respectively, a template ordered as *S. aureus* codon optimized synthetic double stranded DNA (IDT) and plasmid pJ201mNeonGreen (ATUM). Restriction ligation and Gibson assembly cloning was done using enzymes from New England Biolabs according to the manufacturer’s instructions. All cloned constructs were validated by PCR and sequencing. PCR- and gel purification, as well as plasmid isolation, were done using QIAquick PCR purification kit, QIAquick PCR & Gel cleanup kit and QIAprep Spin Miniprep kit (all from Qiagen), according to the manufacturer’s instructions. Cloning of plasmids to be used for *S. aureus* strain generation was done in *E. coli* strain DC10B due to its suitable methylation pattern[50].

### Strain construction

Plasmids were introduced into electrocompetent *S. aureus* RN4220 cells as previously described [51] and transduced to JE2 using phage 80α [52]. Antibiotic marker-free allelic replacements of the *spa* gene by constructs of interest, in the *S. aureus* chromosome, were performed using plasmids that allow for double homologous recombination events at the *spa* locus sites, except for strain BCBNM053 in which the allelic replacement took place at the *SAUSA300_2100* site. A mNG derivative, mNG^ecySA^, which is mNG^Δ1-9^ lacking the first 9 amino acids, optimised for improved expression and N-terminal FP fusion production [53], was used for strains with mNG as an N-terminal fusion to PBP1 (BCBNM031), DivIB (BCBNM037), FtsA (BCBNM032) and ZapA (BCBNM035). For simplicity, mNG is used to describe, mNG^ecySA^ in these strains.

### Cloning, expression and purification of His-sfTq2^ox^ and His-mTq2

Plasmids, pET-sfTq2^ox^ and pET-mTq2 were introduced into *E. coli* BL21 (DE3) for expression of His_6_-sfTq2^ox^ and His_6_-mTq2, respectively. Transformants were grown at 37 °C in LB medium containing 50 μg/ml Kanamycin. When OD_600_ values were ∼0.6 - 0.8, the cultures were supplemented with 1 mM IPTG and grown for an additional 3 hours. Cells were harvested by centrifugation, resuspended in Buffer A (50 mM Sodium Phosphate buffer (pH 8) containing 150 mM NaCl and complete-EDTA-free protease inhibitors, Roche) and were disrupted by sonication (Branson, 40% amplitude, 10 times 10 seconds in an ice/water bath with 1 minute cooling in between cycles). After centrifugation at 48,000 g at 4 °C, the soluble fraction was applied to pre-equilibrated Talon™ (Clontech) resin and incubated at 4 °C overnight. Bound protein was washed twice with Buffer A, and then once with Buffer A containing 10 mM imidazole. Bound protein was eluted in a two-step manner with Buffer A containing 100 mM then 150 mM imidazole. Eluted fractions were combined and dialysed sequentially against 50 mM sodium phosphate buffer (pH 8) containing 500 mM NaCl, 300 mM NaCl and 150 mM NaCl. The concentration of the purified proteins (6 mg / ml for His_6_-sfTq2^ox^ and 5 mg / ml for His_6_-mTq2) was quantified using the Pierce BCA Protein Assay kit (Thermo Scientific) and purity was estimated to be at least 95 % by visual inspection of Coomassie stained SDS-PAGE.

### FLIM experiments

Precultures of *S. aureus* strains were made inoculating a single colony in 5 ml EZ-rich (which is supplemented with 5% TSB) and grown ∼16-18h at 30 °C while shaking at 200 rpm. The overnight cultures were diluted 1:1000-2000 into 20 ml of fresh medium and cultured at 30 °C while shaking. If required, production of fusion proteins was induced with 2-50 ng/ml of anhydrous tetracycline (aTc, Sigma) at OD_600_ ∼0.025-0.05. Cultures were grown to OD_600_ ∼0.3-0.6 (log phase) and concentrated by centrifugation of the culture for 10 min at 7000 rpm. The pellet was washed with PBS and concentrated for 5 min at 7000 rpm. The supernatant was discarded and the cells resuspended in the leftover volume. A volume of 2.5 µl of the cell suspension was spotted on agarose (Biorad #1613100) pad (1.2 % w/v in PBS, containing 0.4 % glucose) in a gene frame (ABgene) on a microscopy slide (VWR) and covered with a No. 1.5 High performance coverslip (Zeiss).

For the FLIM-FRET experiments with FtsW-HT, overnight cultures were prepared as described above. The overnight cultures were diluted 1:2000 in 20 ml of fresh medium and continued to grow at 30 °C. At OD_600_ ∼0.025, FtsW-HT production was induced with 2 ng / ml aTc and at OD_600_ ∼0.4-0.5, 15 ml culture was concentrated in 1 ml fresh medium containing 25 nM of the JF503 dye or 25 nM of a 1:1 mix of JF503 and JF585 dyes, and incubated at 30 °C, shaking at 900 rpm for 30 min. The cells were then washed twice with EZ-rich medium before microscopy slides were prepared as described above.

### Imaging conditions

Fluorescence lifetime images were acquired using a Zeiss LSM880 confocal microscope (Carl Zeiss Microscopy GmbH, Germany) equipped with a FLIM upgrade kit consisting of a Time Harp 260 Pico dual TCSPC Unit, 2 PMA-Hybrid 40 detectors and an LDH-D-C440 pulsed diode laser controlled by a PDL 800-d dual mode driver (Picoquant GmbH, Germany). Microscopy slides were mounted on the stage and a field of view of 440×440 pixels was scanned with 440-nm laser at 20 MHz (Max. optical power 2.2 mW, measured at the collimator’s aperture, attenuated to <10 %) through a H437/25 (AFH GmbH, Germany) clean up filter and a Zeiss MBS445 filter using a Zeiss objective lens Plan-APOCHROMAT QS 63x oil DIC numeric aperture 1.4. Fluorescence signal was detected using pinhole set between 100-200 µm. For sfTq2^ox^-mNG FLIM-FRET experiments, the fluorescence signal was split into donor and acceptor light paths by a H488LPXP (AFH) dichroic mirror. Donor and acceptor emission signals were respectively detected through a FF01-470/28 and a FF01-550/49 filter (Semrock, USA). For FLIM-FRET experiments using JF503 and JF585 dyes, fluorescence signal was split into donor and acceptor light paths by a FF580-FDi01 dichroic mirror (Semrock) and the fluorescence signals were respectively detected through a FF01-520/44 and a FF01-630/92 filter (Semrock). Photon arrival times were collected by time correlated single photon counting (TCSPC). The resulting fluorescence lifetime images were screened by visual inspection for obvious outliers, such as dead cells or cells that were disproportionally bright, which were then excluded from subsequent analysis and appear greyed-out in presented images. Amplitude weighted fluorescence lifetimes were obtained by fitting the fluorescence decay curves using a three-exponential reconvolution model incorporating an instrument response function (IRF) generated from the same data set. The scatter and/or reflection component (<100 ps) was later excluded for calculating the average fluorescence lifetime using SymPhoTime 64 v2.6 and/or v2.8 (Picoquant) software (**Supplementary Text 3**). Fitted results were accepted or discarded based on visual inspection of the fit, the residuals and minimal χ^2^ values. The reported fluorescence lifetimes are amplitude weighted averages (τ_amp_). The FastFLIM (or Fast Lifetime) images shown represent the mean arrival time of photons after the laser pulse. In detail, Fast Lifetime is calculated based on the barycentre of the pixel’s decay. The timespan from the barycentre of the IRF to the barycentre of the decay equals the average lifetime (Picoquant). As a reference for instrument performance, the fluorescence lifetime Alexa 488 (5 µg/ml) was measured periodically using the same excitation/emission settings applied to sfTq2^ox^. The average fluorescence lifetime of this dye was obtained by one-component reconvolution fitting.

### Spectral-based FRET

Spectral-based FRET experiments were performed as previously described [23,31]. In short, spectral unmixing dissects the fluorescence spectra measured for strains with fluorescent proteins undergoing FRET into its individual spectra components. This includes donor, acceptor and background fluorescence. When FRET occurs, there is also a fourth spectral component with the same spectrum as the acceptor, but that cannot be accounted for based on reference spectra. This is called sensitized emission and is used to calculate acceptor-based FRET efficiency (E*f*A). *S. aureus* strain JE2, or its derivatives expressing sfTq2^ox^ or mNG were used as references for background, sfTq2^ox^ and mNG spectra, respectively. These reference strains, as well as the strain expressing protein fusions whose interaction is being assessed, were grown and fusion protein production was induced as described above for the FLIM experiments. The cells were then harvested, washed with PBS and OD_450_ was adjusted to values of 1.000 ± 0.005. Replicate samples of 200 µl (4 to 8, based on the available volume) were loaded in clear-bottom black-walled polystyrene 96-well plate (Corning 3606). Fluorescence spectra were measured in a Neo2 multimode plate reader (Biotek) with excitation at 450 nm and emission between 470-650 nm for the donor sfTq2^ox^ and with 495 nm excitation and emission between 517-650 nm for the acceptor mNG. The fluorescence spectra of the replicates were averaged, the resulting references and sample spectra were unmixed into their background, donor, acceptor and sensitized emission spectra and the acceptor-based FRET efficiencies (E*f*A) were calculated as described previously [23,31].

## Acknowledgements

We are grateful for the help of Mark Hink (University of Amsterdam) and Volker Bushmann (Picoquant, Berlin) in setting up the data analysis pipeline. Additionally, we are grateful for the suggestions by Tanneke den Blaauwen (University of Amsterdam) after critically reading the manuscript. We thank Luke Lavis (Janelia Research Campus, Ashburn) for the generous gift of JF503-HTL and JF585-HTL. This study was funded by the European Research Council (ERC) through grants ERC-2017-CoG-771709 and ERC-2022-ADG 101096393 (to M.G.P.); by Fundação para a Ciência e a Tecnologia (FCT) through MOSTMICRO-ITQB R&D Unit (UIDB/04612/2020, UIDP/04612/2020 to ITQB-NOVA), LS4FUTURE Associated Laboratory (LA/P/0087/2020 to ITQB-NOVA), Individual Call to Scientific Employment Stimulus (2022.00548.CEECIND to N.Y.M.), Call for Exploratory Projects in All Scientific Domains 2023 (2023.12822.PEX, to N.Y.M.), PhD fellowship PD/BD/135480/2018 (to S.F.C.); by the European Union’s Horizon 2020 research and innovation programme under the Marie Sklodowska-Curie grant agreement Nº 839596 (to S.S.) and by the Portuguese Platform of BioImaging (PPBI-POCI-01-0145-FEDER-022122). FF was supported by UIDB/04565/2020 and UIDP/04565/2020 of the Research Unit iBB-Institute for Bioengineering and Biosciences, and the project LA/P/0140/2020 of the Associate Laboratory i4HB-Institute for Health and Bioeconomy.

## References

1. Du, S. & Lutkenhaus, J. Assembly and activation of the Escherichia coli divisome. Mol. Microbiol. 105, 177– 187 (2017).

2. Adams, D. W. & Errington, J. Bacterial cell division: Assembly, maintenance and disassembly of the Z ring. Nat. Rev. Microbiol. 7, 642–653 (2009).

3. Pinho, M. G. & Foster, S. J. Cell Growth and Division of Staphylococcus aureus. Annu. Rev. Microbiol. 78, 293–310 (2024).

4. Pichoff, S. & Lutkenhaus, J. Tethering the Z ring to the membrane through a conserved membrane targeting sequence in FtsA. Mol. Microbiol. 55, 1722–1734 (2005).

5. Fujita, J. et al. Crystal structure of FtsA from Staphylococcus aureus. FEBS Lett. 588, 1879–1885 (2014).

6. Bisson-Filho, A. W. et al. Treadmilling by FtsZ filaments drives peptidoglycan synthesis and bacterial cell division. Science 355, 739–743 (2017).

7. Yang, X. et al. GTPase activity-coupled treadmilling of the bacterial tubulin FtsZ organizes septal cell wall synthesis. Science. 355, 744–747 (2017).

8. Gamba, P., Veening, J. W., Saunders, N. J., Hamoen, L. W. & Daniel, R. A. Two-step assembly dynamics of the Bacillus subtilis divisome. J. Bacteriol. 191, 4186–4194 (2009).

9. Tinajero-Trejo, M. et al. Control of morphogenesis during the Staphylococcus aureus cell cycle. Sci. Adv. 11, 1–16 (2025).

10. Monteiro, J. M. et al. Peptidoglycan synthesis drives an FtsZ-treadmilling-independent step of cytokinesis. Nature 554, 528–532 (2018).

11. Aarsman, M. E. G. et al. Maturation of the Escherichia coli divisome occurs in two steps. Mol. Microbiol. 55, 1631–1645 (2005).

12. Reichmann, N. T. et al. SEDS–bPBP pairs direct lateral and septal peptidoglycan synthesis in Staphylococcus aureus. Nat. Microbiol. 4, 1368–1377 (2019).

13. Taguchi, A. et al. FtsW is a peptidoglycan polymerase that is functional only in complex with its cognate penicillin-binding protein. Nat. Microbiol. 4, 587–594 (2019).

14. Schäper, S. et al. Cell constriction requires processive septal peptidoglycan synthase movement independent of FtsZ treadmilling in Staphylococcus aureus. Nat. Microbiol. 9, 1049–1063 (2024).

15. Pazos, M. et al. Z-ring membrane anchors associate with cell wall synthases to initiate bacterial cell division. Nat. Commun. 9, 5090 (2018).

16. Söderström, B., Chan, H., Shilling, P. J., Skoglund, U. & Daley, D. O. Spatial separation of FtsZ and FtsN during cell division. Mol. Microbiol. 107, 387–401 (2018).

17. Lyu, Z. et al. FtsN maintains active septal cell wall synthesis by forming a processive complex with the septum-specific peptidoglycan synthases in E. coli. Nat. Commun. 13, 5751 (2022).

18. Suhling, K., French, P. M. W. & Phillips, D. Time-resolved fluorescence microscopy. Photochem. Photobiol. Sci. 4, 13–22 (2005).

19. Gadella, T. W. J. FRET and FLIM techniques. (Elsevier, 2009).

20. Alexeeva, S., Gadella, T. W. J., Verheul, J., Verhoeven, G. S. & den Blaauwen, T. Direct interactions of early and late assembling division proteins in Escherichia coli cells resolved by FRET. Mol. Microbiol. 77, 384–98 (2010).

21. Meiresonne, N. Y., van der Ploeg, R., Hink, M. A. & den Blaauwen, T. Activity-related conformational changes in D,D-carboxypeptidases revealed by in vivo periplasmic förster resonance energy transfer assay in Escherichia coli. MBio 8, e01089–17 (2017).

22. Meiresonne, N. Y. & Den, T. The in vitro non-tetramerizing ZapAI83E mutant is unable to recruit ZapB to the division plane in vivo in Escherichia coli. Int. J. Mol. Sci. 21, 3130 (2020).

23. Meiresonne, N. Y. et al. Superfolder mTurquoise2 ox optimized for the bacterial periplasm allows high efficiency in vivo FRET of cell division antibiotic targets. Mol. Microbiol. 111, 1025–1038 (2019).

24. Manko, H. et al. PvdL Orchestrates the Assembly of the Nonribosomal Peptide Synthetases Involved in Pyoverdine Biosynthesis in Pseudomonas aeruginosa. Int. J. Mol. Sci. 25, 6013 (2024).

25. Manko, H. et al. FLIM-FRET Measurements of Protein-Protein Interactions in Live Bacteria. J. Vis. Exp. (2020). doi:10.3791/61602-v

26. Detert Oude Weme, R.G.J. et al. Single cell FRET analysis for the identification of optimal FRET-Pairs in Bacillus subtilis using a prototype MEM-FLIM system. PLoS One 10, (2015).

27. Mastop, M. et al. Characterization of a spectrally diverse set of fluorescent proteins as FRET acceptors for mTurquoise2. Sci. Rep. 7, 11999 (2017).

28. Pereira, P. M., Veiga, H., Jorge, A. M. & Pinho, M. G. Fluorescent reporters for studies of cellular localization of proteins in Staphylococcus aureus. Appl. Environ. Microbiol. 76, 4346–4353 (2010).

29. Sillen, A. & Engelborghs, Y. The Correct Use of ‘Average’ Fluorescence Parameters. Photochem. Photobiol. 67, 475–486 (1998).

30. Clegg, R. M. Fluorescence resonance energy transfer and nucleic acids. Methods Enzymol. 211, 353–88 (1992).

31. Meiresonne, N., Consoli, E., Mertens, L. & den Blaauwen, T. Detection of in vivo Protein Interactions in All Bacterial Compartments by Förster Resonance Energy Transfer with the Superfolder mTurquoise2 ox-mNeongreen FRET Pair. BIO-PROTOCOL 9, (2019).

32. Karimova, G., Pidoux, J., Ullmann, A. & Ladant, D. A bacterial two-hybrid system based on a reconstituted signal transduction pathway. Proc. Natl. Acad. Sci. U. S. A. 95, 5752–6 (1998).

33. Karimova, G., Dautin, N. & Ladant, D. Interaction network among Escherichia coli membrane proteins involved in cell division as revealed by bacterial two-hybrid analysis. J. Bacteriol. 187, 2233–43 (2005).

34. Käshammer, L. et al. Cryo-EM structure of the bacterial divisome core complex and antibiotic target FtsWIQBL. Nat. Microbiol. 8, 1149–1159 (2023).

35. Park, K. T., Pichoff, S., Du, S. & Lutkenhaus, J. FtsA acts through FtsW to promote cell wall synthesis during cell division in Escherichia coli. Proc. Natl. Acad. Sci. U. S. A. 118, e2107210118 (2021).

36. Liu, B., Persons, L., Lee, L. & de Boer, P. A. J. Roles for both FtsA and the FtsBLQ subcomplex in FtsN-stimulated cell constriction in Escherichia coli. Mol. Microbiol. 95, 945–970 (2015).

37. Steele, V. R., Bottomley, A. L., Garcia-Lara, J., Kasturiarachchi, J. & Foster, S. J. Multiple essential roles for EzrA in cell division of Staphylococcus aureus. Mol. Microbiol. 80, 542–555 (2011).

38. Bertsche, U., Breukink, E., Kast, T. & Vollmer, W. In vitro murein (peptidoglycan) synthesis by dimers of the bifunctional transglycosylase-transpeptidase PBP1B from Escherichia coli. J. Biol. Chem. 280, 38096–38101 (2005).

39. Bon, C. G. et al. Structural and kinetic analysis of the monofunctional Staphylococcus aureus PBP1. J. Struct. Biol. 216, 108086 (2024).

40. Choi, Y. et al. Structural Insights into the FtsQ/FtsB/FtsL Complex, a Key Component of the Divisome. Sci. Rep. 8, 18061 (2018).

41. Condon, S. G. F. et al. The FtsLB subcomplex of the bacterial divisome is a tetramer with an uninterrupted FtsL helix linking the transmembrane and periplasmic regions. J. Biol. Chem. 293, 1623–1641 (2018).

42. Nguyen, H. T. V. et al. Structure of the heterotrimeric membrane protein complex FtsB-FtsL-FtsQ of the bacterial divisome. Nat. Commun. 14, 1903 (2023).

43. Grimm, J. B. et al. A general method to fine-tune fluorophores for live-cell and in vivo imaging. Nat. Methods 14, 987–994 (2017).

44. Los, G. V. et al. HaloTag: A Novel Protein Labeling Technology for Cell Imaging and Protein Analysis. ACS Chem. Biol. 3, 373–382 (2008).

45. Yang, Y., Bhachech, N. & Bush, K. Biochemical comparison of imipenem, meropenem and biapenem: Permeability, binding to penicillin-binding proteins, and stability to hydrolysis by β-lactamases. J. Antimicrob. Chemother. 35, 75–84 (1995).

46. Kuk, A. C. Y., Hao, A. & Lee, S.-Y. Structure and Mechanism of the Lipid Flippase MurJ. Annu. Rev. Biochem. 91, 705–729 (2022).

47. Huber, J. et al. Chemical Genetic Identification of Peptidoglycan Inhibitors Potentiating Carbapenem Activity against Methicillin-Resistant Staphylococcus aureus. Chem. Biol. 16, 837–848 (2009).

48. Holtje, J.-V. A hypothetical holoenzyme involved in the replication of the murein sacculus of Escherichia coli. Microbiology 142, 1911–1918 (1996).

49. Höltje, J. V. Growth of the stress-bearing and shape-maintaining murein sacculus of Escherichia coli. Microbiol. Mol. Biol. Rev. 62, 181–203 (1998).

50. Monk, I.R.,, Shah, I.M.,, Xu, M.,, Tan, M.W.,, Foster, T.J. Transforming the Untransformable: Application of Direct Transformation to Manipulate Genetically Staphylococcus aureus and Staphylococcus epidermidis. mBio, 3:e00277–11 (2012)

51. Veiga, H. & Pinho, M. G. Inactivation of the saul type I restriction-modification system is not sufficient to generate Staphylococcus aureus strains capable of efficiently accepting foreign DNA. Appl. Environ. Microbiol. 75, 3034–3038 (2009).

52. Oshida, T. & Tomasz, A. Isolation and characterization of a Tn551-autolysis mutant of Staphylococcus aureus. J. Bacteriol. 174, 4952–4959 (1992).

53. Mertens, L. M. Y. & den Blaauwen, T. Optimising expression of the large dynamic range FRET pair mNeonGreen and superfolder mTurquoise2ox for use in the Escherichia coli cytoplasm. Sci. Rep. 12, 17977 (2022).

