## Supplementary material for "FtsW protein-protein interactions visualized in live *Staphylococcus aureus* cells by FLIM-FRET"

### Supplementary text 1: Validation of sfTq2$^{ox}$ using time-correlated single photon counting (TCSPC).

To set up FLIM-FRET as a robust technique to demonstrate protein interactions in *Staphylococcus aureus*, a fluorescent protein FRET-pair was chosen with favourable spectroscopic properties. Superfolder mTurquoise2ox (sfTq2$^{ox}$) is a variant of the fluorescent protein (FP) mTurquoise2 sharing its long, monoexponentially decaying, fluorescence lifetime of ~4 ns and high quantum yield of 93 % making it an excellent FLIM-FRET donor [1–3]. Paired with the spectrally compatible acceptor FP mNeonGreen (mNG, EC=116,000 $M^{-1}$ $cm^{-1}$), it forms a FRET-pair with a Förster radius $R_0$ of 6.0 nm [4], which is a substantial (theoretical) improvement over other FRET-pairs previously used in bacteria, such as sfGFP-mCherry ($R_0$ = 5.2 nm) or mNG-mCherry ($R_0$ = 5.5 nm) [5,6]. Indeed, the sfTq2$^{ox}$-mNG FRET pair has been shown to enable detection of energy transfer up to 40 % for periplasmic control proteins in *E. coli,* as demonstrated by spectral unmixing [1].

The spectroscopic properties of sfTq2$^{ox}$ were shown to be identical to those of mTq2, mostly by frequency domain fluorescence lifetime measurements. In this work we use time domain fluorescence lifetime measurements by time-correlated single photon counting (TCSPC). To assess the suitability of sfTq2$^{ox}$, the protein was isolated for spectroscopic studies and compared with mTq2 protein. First, their absorption and fluorescence emission spectra were measured and confirmed to match (**Figure S1a**). Then, their fluorescence decay curves were measured using TCSPC. Both proteins exhibited matching monoexponential decays, which were fitted using one-component reconvolution fitting, yielding a fluorescence lifetime (τ) of ~4.1 ns. (**Figure S1b**).

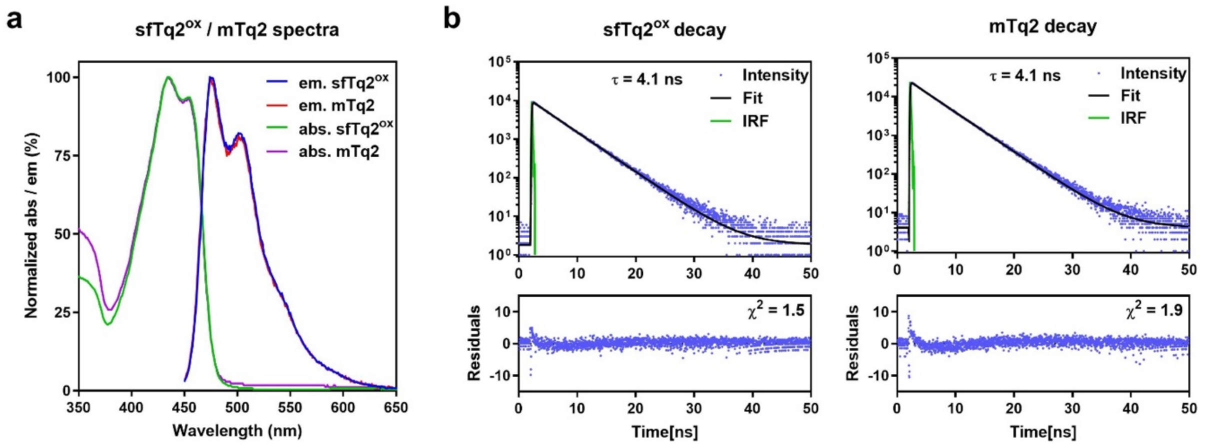

#### Figure S1 – Fluorescence spectra and fluorescence decay curves of sfTq2$^{ox}$ protein

**a**) Normalized absorption and emission spectra of isolated sfTq2$^{ox}$. Isolated mTq2 was used for comparison showing that spectra for the two proteins are very similar. em = emission, abs = absorbance **b**) Fluorescence decay curves of sfTq2$^{ox}$ and mTq2 are similar and yielded the same fluorescence lifetime. τ = fluorescence lifetime, IRF = Instrument response function.

#### Supplementary text 2: Assessment of different growth media for FLIM-FRET assays in *S. aureus*

Culturing conditions were optimized to minimize autofluorescence potentially impairing lifetime measurements in living *S. aureus* cells. Autofluorescence of tryptic soy broth (TSB), the culture medium in which *S. aureus* is routinely grown, was initially assessed. Exciting TSB at the wavelength used for sfTq2$^{ox}$ excitation resulted in a strong emission curve overlapping with the emission of sfTq2$^{ox}$ (**Figure S2a**). This suggested that background/autofluorescence from this medium could become part of the signal in the FLIM measurements and prompted the search for a low-fluorescent growth medium. The often used, and relatively clearer, M9 minimal medium (M9) was tested and showed less fluorescence (**Figure S2a**). However, *S. aureus* cells grow slowly in M9 (**Figure S2b**). We then tested EZ-rich defined medium, a rich medium that is clear and often used to grow Gram-negative bacteria for experiments that require low autofluorescence. Exciting EZ-rich at wavelengths used for sfTq2$^{ox}$ resulted in a relatively weak emission spectrum compared to TSB (25 x lower) and even M9 (3x lower). However, we observed that dilutions of *S. aureus* overnight cultures equal or higher than 1:1000 into fresh EZ-rich led to impaired growth. This led to the hypothesis that EZ-rich may miss one or more components for ideal *S. aureus* growth and supplementation with TSB was tested. *S. aureus* JE2 was grown at 30°C in TSB or EZ supplemented with 5 % TSB to $OD_{600}$ ~0.4 and diluted to 0.002 into fresh media with a respective concentration range from 100/0 % to 50/50 % of EZ-rich/TSB. The cells were then incubated in a plate reader at 30 °C while shaking, with $OD_{600}$ measurements every 15 minutes (**Figure S2c,d)**. Supplementing EZ-rich with only 1 % TSB resulted in almost similar growth compared with the highest concentration of TSB supplementation. For FLIM-FRET experiments, we selected a medium composed of 95 % EZ-rich and 5 % TSB as this was the lowest concentration of TSB that resulted in growth virtually indistinguishable from the highest concentration of TSB tested. For simplicity, EZ Defined Rich supplemented with 5 % TSB will be named "EZ-rich" in the subsequent sections.

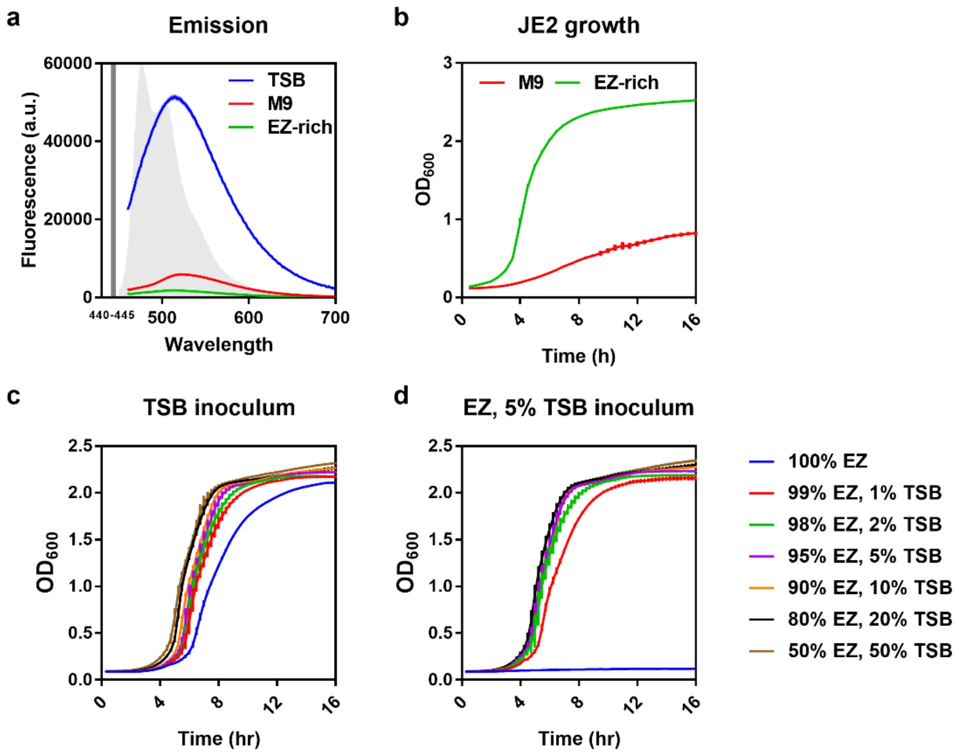

#### Figure S2 – Assessment of auto fluorescence of growth media for FLIM-FRET assays in *S. aureus*

**a**) Emission spectra of Tryptic Soy Broth (TSB), M9 Minimal Medium (M9) or EZ-rich Defined Medium (EZ) upon excitation at 440-445 nm (dark grey line). The light grey curve indicates the emission spectrum of sfTq2$^{ox}$. **b**) Growth curves of *S. aureus* JE2 grown in M9 or EZ-rich medium at 30 °C. **c,d**) Growth curves of *S. aureus* JE2 grown at 30 °C in different ratios of EZ-rich and TSB using an inoculum grown in TSB (c) or in EZ-rich medium with 5% TSB (d). Error bars represent the standard deviation.

#### Supplementary text 3 – Analysis used for fitting fluorescence decay curves and removal of scattering signal

Purified sfTq2$^{ox}$ protein exhibited a fluorescence lifetime of 4.1 ns, as described in **Supplementary text 1**. Measuring the lifetime of sfTq2$^{ox}$ in the biological context of a *S. aureus* cell requires considering its local environment. Indeed, lower lifetimes are often measured *in vivo,* compared to the lifetime of the isolated fluorescent protein [2,3,5]. Therefore, fitting of the fluorescence decay curves is done with two components to accommodate for the fluorescent protein as well as the cellular environment. During initial experiments we noticed that fluorescence lifetime measurements of *S. aureus* producing sfTq2$^{ox}$ could result in two different values, with amplitude weighted averages either (i) approaching the expected value of 4.0 ns, using two-component fitting or (ii) having extremely low values, close to 0.0 ns. In the latter cases, a 3-component fit (R3) revealed individual $\tau_1$ and $\tau_2$ values to be within the expected fluorescent lifetime range (for sfTq2$^{ox}$ and cellular auto fluorescence) with comparable amplitudes. However, the $\tau_3$ lifetime was close to 0.0 ns with an extremely high amplitude (**Figure S3a, b, c**), causing $\tau_{amp}$ values to be severely underestimated and leading us to investigate its cause.

Due to the small size of *S. aureus* cells, imaging microscopy is conducted very close to the coverslip. We hypothesized that the extremely short lifetimes observed were caused by scattering of the excitation light, with the scattered light contributing disproportionately to the measured signal, leading to inaccurate $\tau_{amp}$ values for sfTq2$^{ox}$ in the cells. To address this, we applied a scatter reduction method (which we will refer to as R3*) by fitting the measured fluorescence decay curves with an additional component dedicated to the scatter signal, which was then subtracted for the $\tau_{amp}$ calculations. This approach resulted in average fluorescence lifetimes in better agreement with the expected values for sfTq2$^{ox}$ in cells (**Figure S3a**).

To further investigate the possibility of light scattering at the glass interface, isolated sfTq2$^{ox}$ was imaged at various positions relative this interface (determined using the Zeiss LSM 880 "definite focus find surface" tool, which is often used to determine the imaging plane). The corresponding fluorescence decay curves were measured and fitted, revealing that when the confocal volume is positioned inside the glass, measured sfTq2$^{ox}$ fluorescence lifetimes were lower than expected. From the interface onwards, the lifetimes increased and reached the expected lifetime of sfTq2$^{ox}$ when the confocal volume is fully inside the protein suspension (**Figure S3d**). This effect was also visible when comparing the normalized fluorescence decay curves of these measurements, which are shorter when the confocal volume is positioned just below the interface or at the interface, than when it is positioned above the coverslip, within the sfTq2$^{ox}$ suspension (**Figure S3e**). These results strongly favour the hypothesis that light scattering at the coverslip glass interface influences the calculated $\tau_{amp}$ values.

To test if the contribution of light scattering could be removed as described above (R3*), in samples with live *S. aureus* cells, strain COL harbouring plasmid pBCBSMC017 (pCNX-sfTq2$^{ox}$-TM) was grown in EZ-rich at 30 °C and production of sfTq2$^{ox}$-TM was induced with 0.1 µM $CdCl_2$ for at least 4 mass doublings. The cells were washed 3 times with PBS and placed on a 1.2 % agarose in PBS pad on a microscopy slide and fluorescence decay curves were measured at the interface or slightly further into the sample. A three-component reconvolution fit yielded an the extremely low $\tau_{amp}$ value at the interface (**Figure S3f,** 0), which increased when measurements were made away from the coverslip (**Figure S3f,** +9μm), as suggested by the fluorescence decay curves (**Figure S3g**). Strikingly, leaving out the scatter component from the $\tau_{amp}$ calculation resulted in the lifetime measured at the interface to be close to the $\tau_{amp}$ obtained from the measurement away from the glass interface (**Figure S3f,** 0*). Considering the 1 µm diameter of *S. aureus* cells*,* imaging at the glass interface is essential, as images taken away from the coverslip quickly lose the necessary resolution and focus (**Figure S3h**). The "definite focus find surface" feature used to define the interface works by detecting infra-red reflection from the coverslip, so scattering is expected at this position. Therefore, for this work, cells were imaged by first using this feature to find the approximate location of the interface, then manually adjusting the imaging plane by visual inspection of focal plane and subsequently using the R3* scatter reduction analysis approach to obtain both spatial resolution and good fluorescence lifetime information from samples.

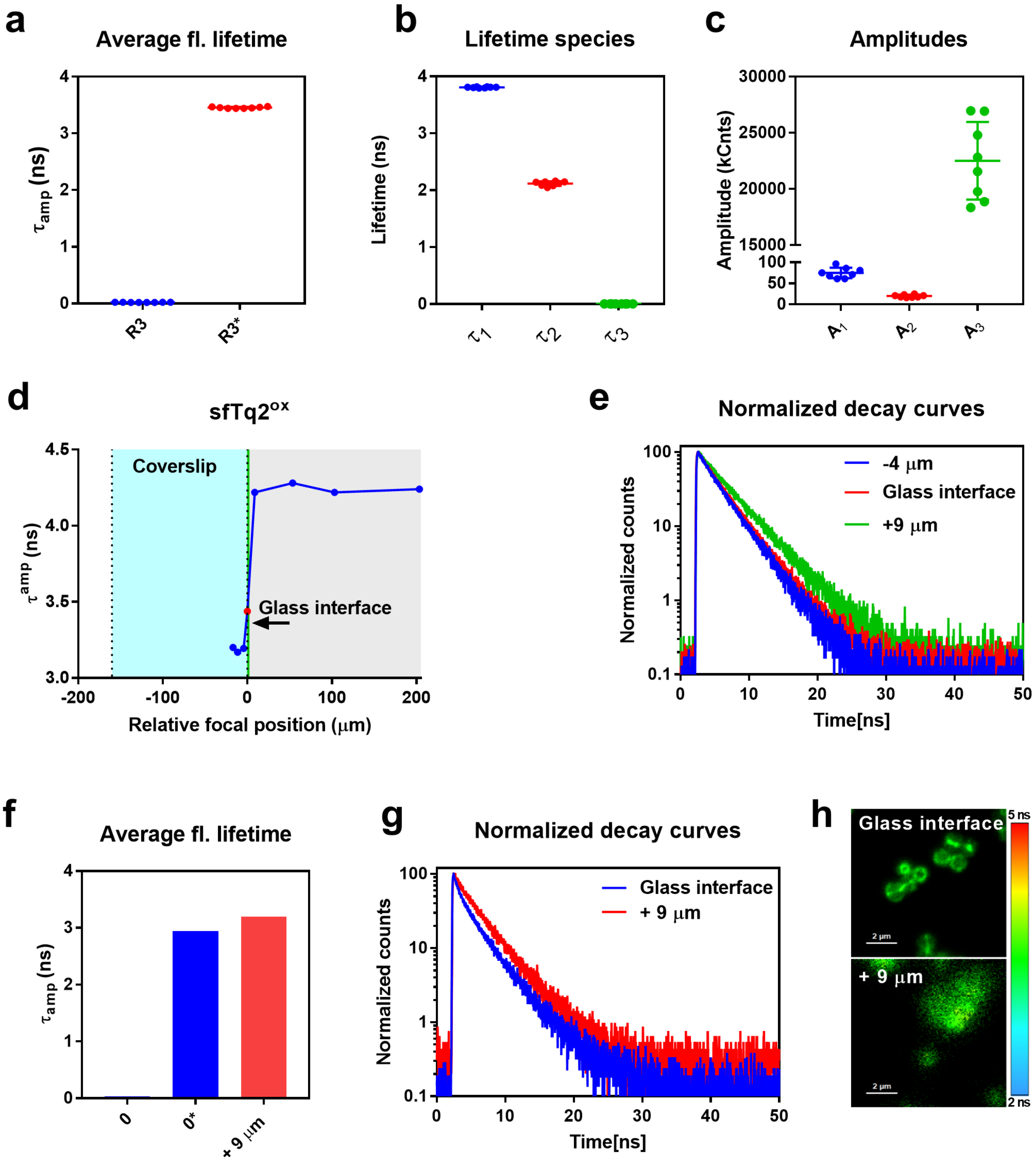

Figure S3 – Assessment of the impact of light scattering at the glass interface on FLIM data.

**a**) Amplitude weighted average fluorescence lifetimes measured from *S. aureus* strain COL producing sfTq2$^{ox}$-TM (from plasmid pBCBSMC017) using a 3-component reconvolution fit (R3) results in extremely low values. Removal of the scattering component recovers this lifetime to expected values (R3*). **b**) Individual lifetime components obtained from the R3 fitting showing an extremely low $\tau_3$ value presumably resulting from light scattering. **c**) Amplitudes associated with the individual fluorescence lifetimes components obtained from the R3 fitting shown in panel b, showing a high $A_3$ value (presumably corresponding to light scattering) relative to $A_1$ and $A_2$. **d**) Amplitude weighted average fluorescence lifetimes of sfTq2$^{ox}$ protein measured with the confocal volume at various positions from the coverslip/sample interface determined by the Zeiss LSM 880 “definite focus find surface” tool. All positions in or close to the glass (blue area) gave lower fluorescence lifetimes, while positions within the protein solution (grey area), away from the glass, gave the expected high lifetimes close to that of sfTq2$^{ox}$. The dashed lines indicate the glass interfaces. **e**) Fluorescence decay curves associated with measurements at the glass interface, above and below (from panel d) showing a longer decay for the measurement in the protein solution, away from the glass. **f**) Average fluorescence lifetimes measured from *S. aureus* COL cells producing sfTq2$^{ox}$-TM from plasmid pBCBSMC017 at the glass interface or away from the glass (different fields of view), showing the scatter-related low lifetime at the glass (0 µm) and a higher lifetime measured away from the glass (+ 9 µm). Removal of the scatter component (0 µm*) resulted in a higher lifetime. **g**) The fluorescence decay curves associated with the 0 µm and + 9 µm measurements (from panel f) showing a longer decay away from the glass. **h**) Fluorescence lifetime images of the data shown in panels f and g.

#### Supplementary text 4: Assessment of bleed-through of mNeonGreen fluorescence into donor detector channel

Potential bleed-through from directly excited acceptor protein mNG into the donor detector channel could theoretically lower the $sfTq2^{ox}$ average lifetime measured in strains where mNG is co-produced with $sfTq2^{ox}$. This concern is based on the fact that the 440 nm laser used to excite the $sfTq2^{ox}$ donor protein, also mildly (~9 % efficiency) excites the acceptor protein mNG ($\tau$ = 3.1 ns) [4]. To minimize the possibility of bleed-through, we used a 488 nm dichroic and 470/28 nm emission filters, given that less than 1 % mNG emission is below 488 nm (**Materials and Methods**). This setup proved to be adequate as we observed similar lifetimes for the donor strain BCBNM002 (producing only $sfTq2^{ox}$) and negative control strain BCBNM005 (producing $sfTq2^{ox}$ and mNG), indicating that measured $sfTq2^{ox}$ lifetimes are not influenced by the presence of non-interacting mNG **(Figure 1**). Furthermore, to fully exclude mNG fluorescence bleed-through as a contributing factor to the measured $sfTq2^{ox}$ lifetimes, strain BCBNM001, constitutively producing only mNG, was constructed and measured under the same conditions as described to measure $sfTq2^{ox}$ lifetimes. This resulted in negligible fluorescence, comparable to the fluorescence measured for the wild-type JE2 cells not producing any fluorescent protein, confirming that the presence of mNG by itself did not affect $sfTq2^{ox}$ lifetime measurements (**Figure S4**).

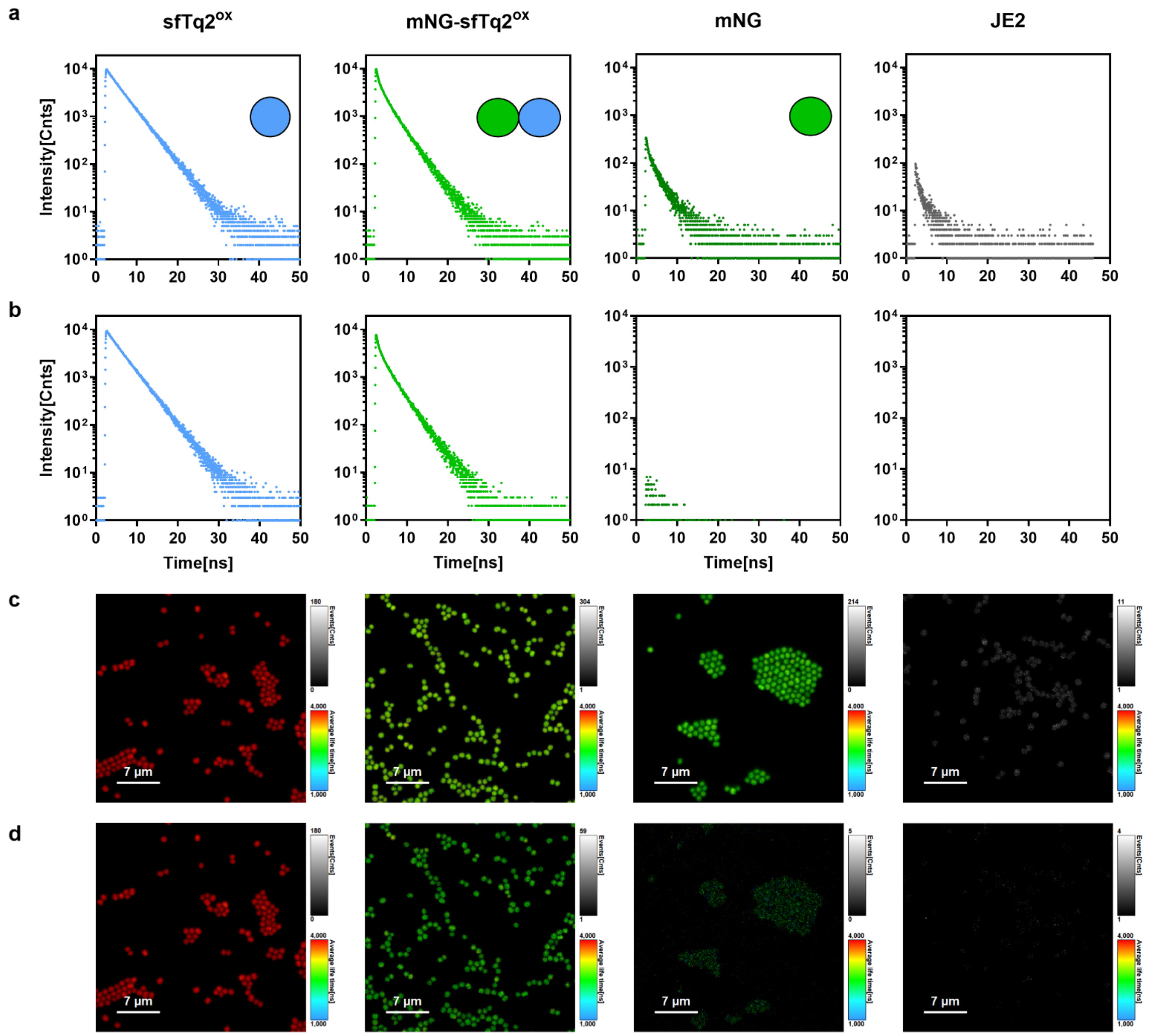

Figure S4 – Assessment of bleed-through of mNeonGreen fluorescence into donor detector channel

**a**) Fluorescence decay curves measured through the sfTq2$^{ox}$ donor channel for JE2 cells producing (from left to right) sfTq2$^{ox}$ (BCBNM002), mNG-sfTq2$^{ox}$ (BCBNM003), mNG (BCBNM001) or no fluorescent protein (JE2). Cartoons represent fluorescent proteins produced in each strain. **b**) Fluorescence decay curves shown in a. after background subtraction. **c**) FastFLIM images (**see Methods**) of the cells corresponding to the fluorescence decay curves shown in a. **d**) FastFLIM images shown in c. after background subtraction.

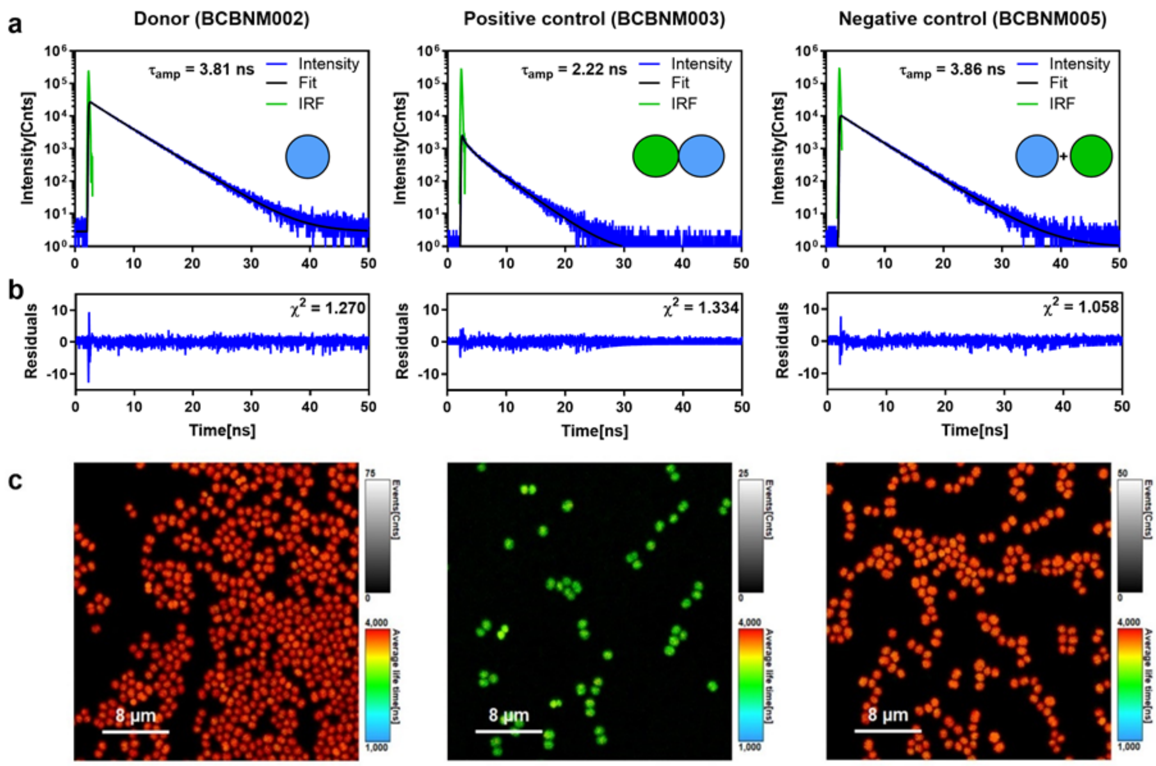

#### Figure S5 – Representative decay curves for the control strains expressing cytosolic fluorescent proteins

**a**) Representative fluorescence decay curves for the indicated control strains expressing cytosolic sfTq2$^{ox}$ (donor), mNG-sfTq2$^{ox}$ (Positive control) and mNG + sfTq2$^{ox}$ (Negative control). Cartoons represent fluorescent proteins produced in each strain. $\tau_{amp}$ = amplitude weighted average fluorescence lifetime, IRF = Instrument Response Function. **b**) Residuals corresponding to the fit of the fluorescence decay curves presented in a. **c**) FastFLIM (**see Methods**) images corresponding to the fluorescence decay curves shown in a.

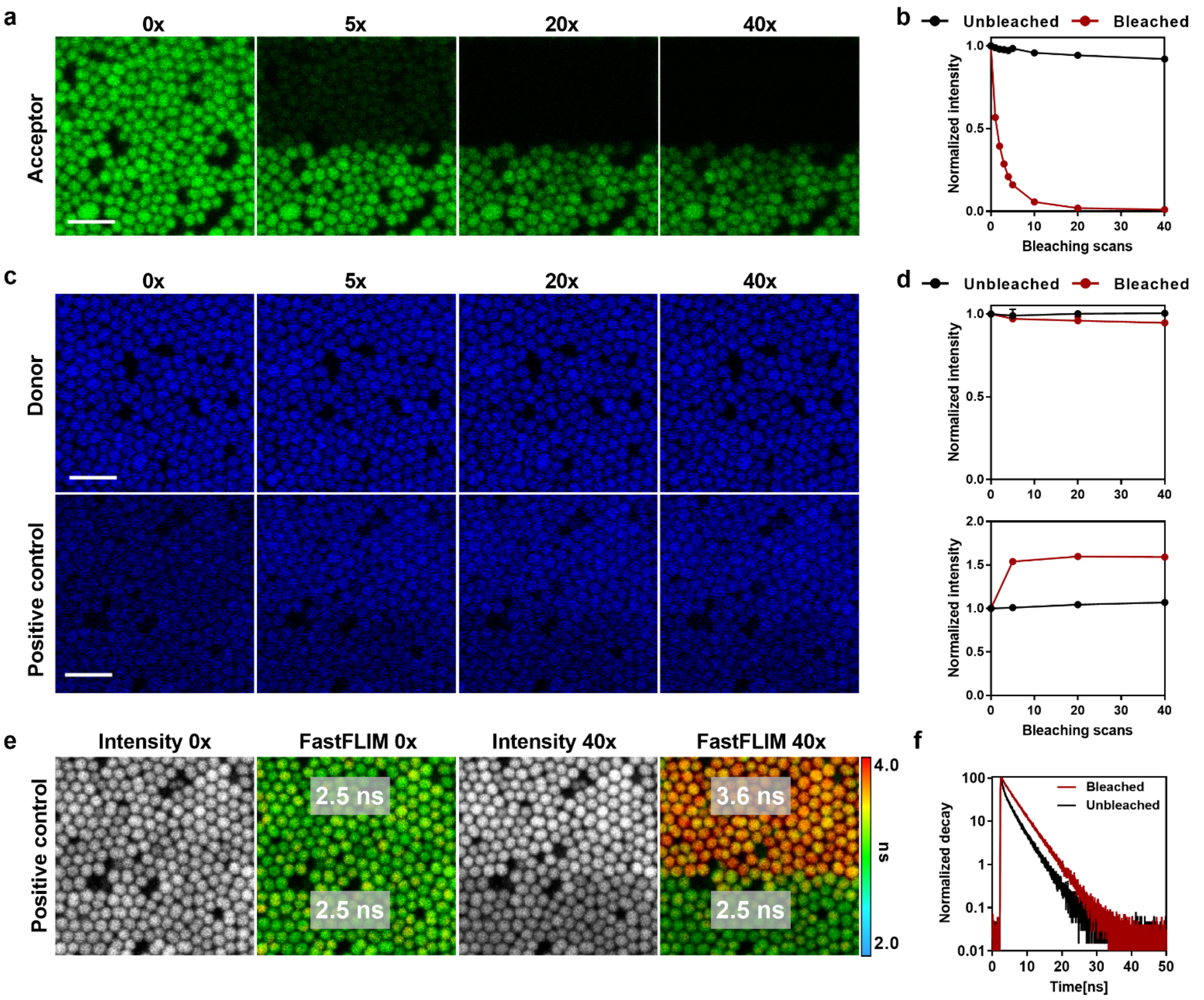

#### Figure S6 – Acceptor photobleaching as a method to validate FLIM-FRET results.

**a**) *S. aureus* JE2 strain producing mNG (Acceptor, BCBNM001) was imaged in the acceptor channel by confocal microscopy before and after bleaching the top region of the field of view with the indicated number of bleaching scans, using the 514 nm laser line at 100 % power. A substantial decrease of mNG signal was observed in the bleached region, with the extent of the signal loss correlating with the number of bleaching scans. In contrast, the unbleached region showed minimal change in intensity. **b**) Quantification of the acceptor fluorescence intensity in bleached and unbleached regions of the images shown in a. **c**) *S. aureus* JE2 strains producing sfTq2$^{ox}$ (Donor, BCBNM002) or the positive control tandem mNG-sfTq2$^{ox}$ (BCBNM003) were imaged in the donor channel by confocal microscopy, before and after bleaching the top region of the field of view with the indicated number of bleaching scans using the 514 nm laser line at 100 % power. As expected, acceptor photobleaching did not affect sfTq2$^{ox}$ fluorescence in the donor strain. In contrast, an increase in sfTq2$^{ox}$ fluorescence intensity was observed in the bleached region of the positive control image, consistent with reduced FRET efficiency upon acceptor photobleaching. **d**) Quantification of the donor fluorescence intensity in acceptor-photobleached and unbleached regions of the images shown in c. **e**) Intensity and FastFLIM (**see Methods**) images of the positive control strain BCBNM003, taken before (two left images) and after (two right images) acceptor photobleaching in the top half of the field of view. Before bleaching both regions exhibit the same average fluorescence lifetime. However, after 40 bleaching scans, the lifetime in the bleached region shifts towards the donor-only lifetime indicating a reduction in FRET efficiency. **f**) Normalized fluorescence decay curves from the bleached and unbleached regions of images of the positive control showing a shift towards a longer lifetime for the bleached region, suggesting strong FRET in the unbleached sample. All scale bars represent 2 µm.

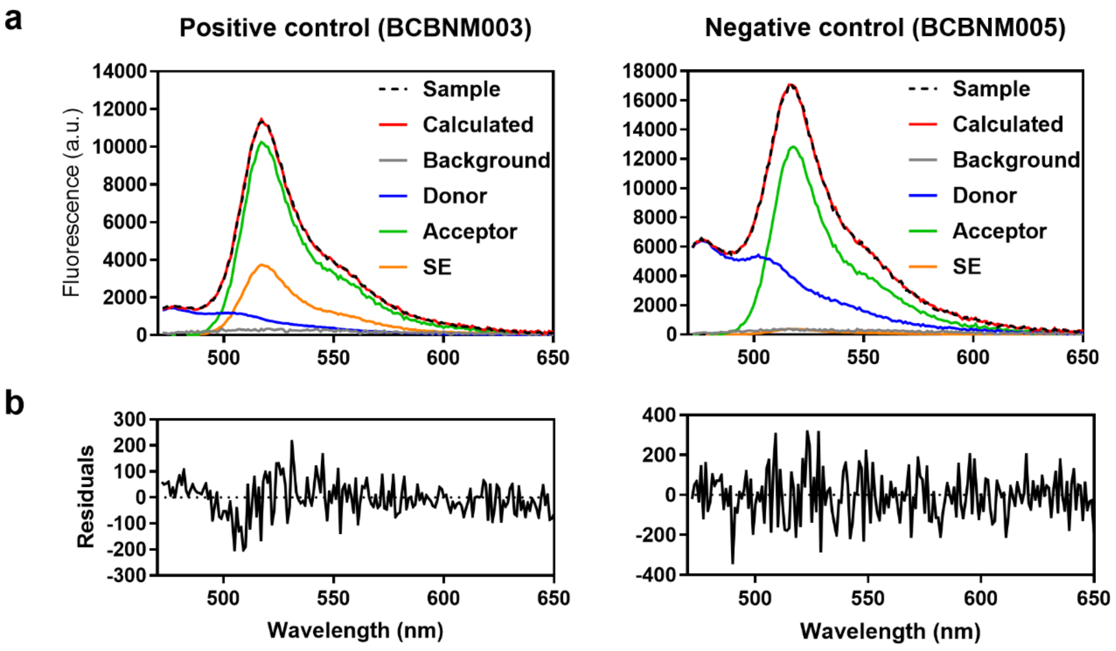

Figure S7 – Representative spectral unmixing graphs of fluorescence spectra measured for the JE2 control strains expressing fluorescent cytosolic proteins.

**a**) Unmixing of the measured acceptor spectra into their individual spectral components to determine acceptor based FRET (**see Methods**) performed as described in Meiresonne *et al* [7]. Reference spectra were obtained from strains BCBNM002 (donor), BCBNM001 (acceptor) and JE2 (background). The black dashed lines show the measured spectrum which is dissected into a spectrum for the background of JE2 cells (gray), sfTq2$^{ox}$ (blue), mNG (green) and the sensitized emission (SE, orange). The sum of the unmixed spectra is represented as a red line (calculated). The resulting E*f*A values were 33.0 ± 2.7 % (n = 7) for the positive control (BCBNM003) and 1.1 ± 0.1 % (n = 5) for the negative control (BCBNM005). **b**) Residual signal that could not be accounted for by unmixing is plotted (measured signal – calculated signal) and served as a measure of quality.

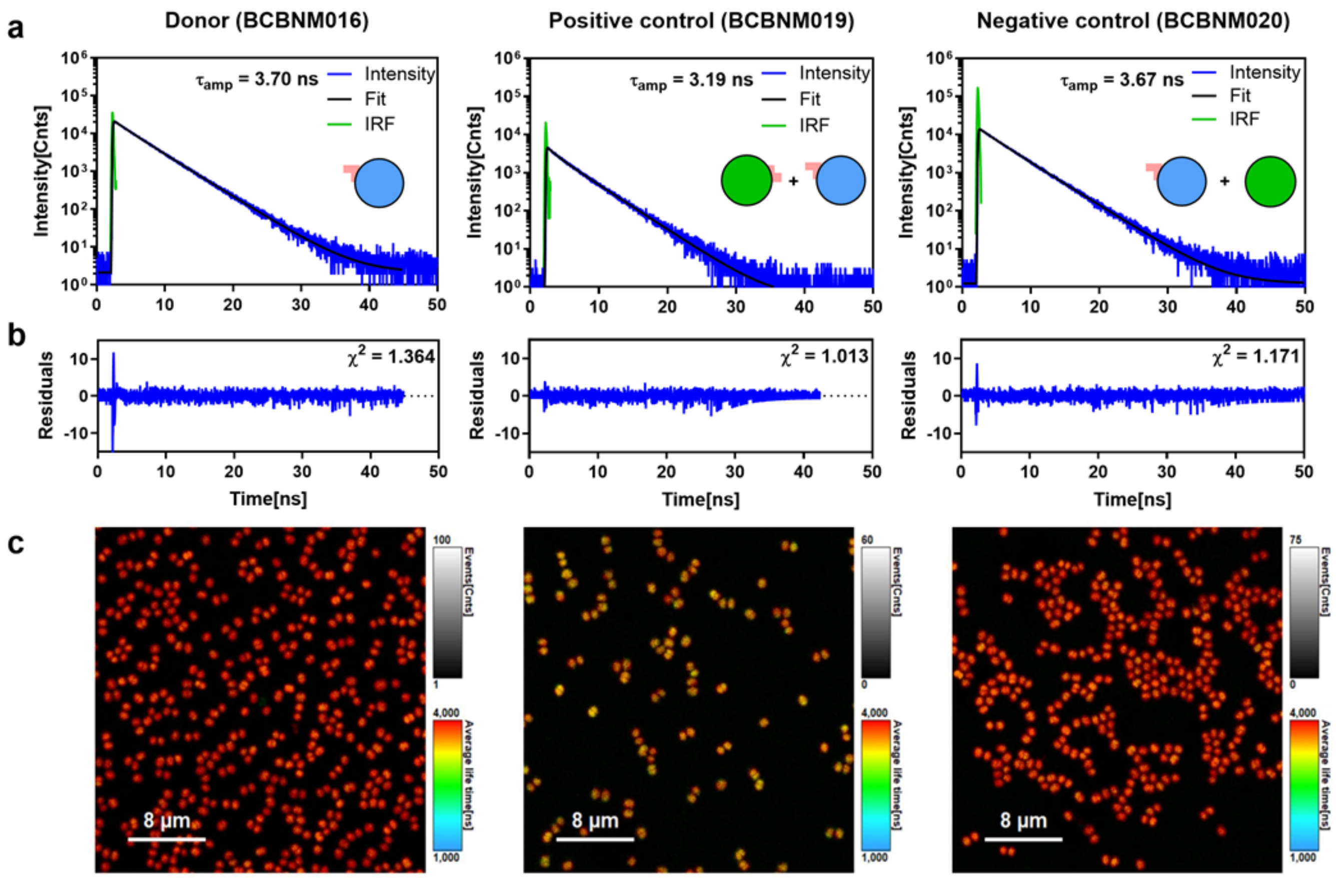

#### Figure S8 – Representative fluorescence decay curves for the Zip-Zip interaction control strains

**a**) Representative fluorescence decay curves for the indicated control strains in JE2 background producing sfTq2$^{ox}$-Zip$^{25}$(Donor), mNG-Zip$^{18}$ + sfTq2$^{ox}$-Zip$^{25}$ (Positive control) and sfTq2$^{ox}$-Zip$^{25}$ + mNG (Negative control). Cartoons represent fluorescent proteins produced in each strain. $\tau_{amp}$ = amplitude weighted average fluorescence lifetime, IRF = Instrument Response Function. **b**) Residuals corresponding to the fit of the fluorescence decay curves presented in a. **c**) FastFLIM (**see Methods**) images corresponding to the fluorescence decay curves shown in a.

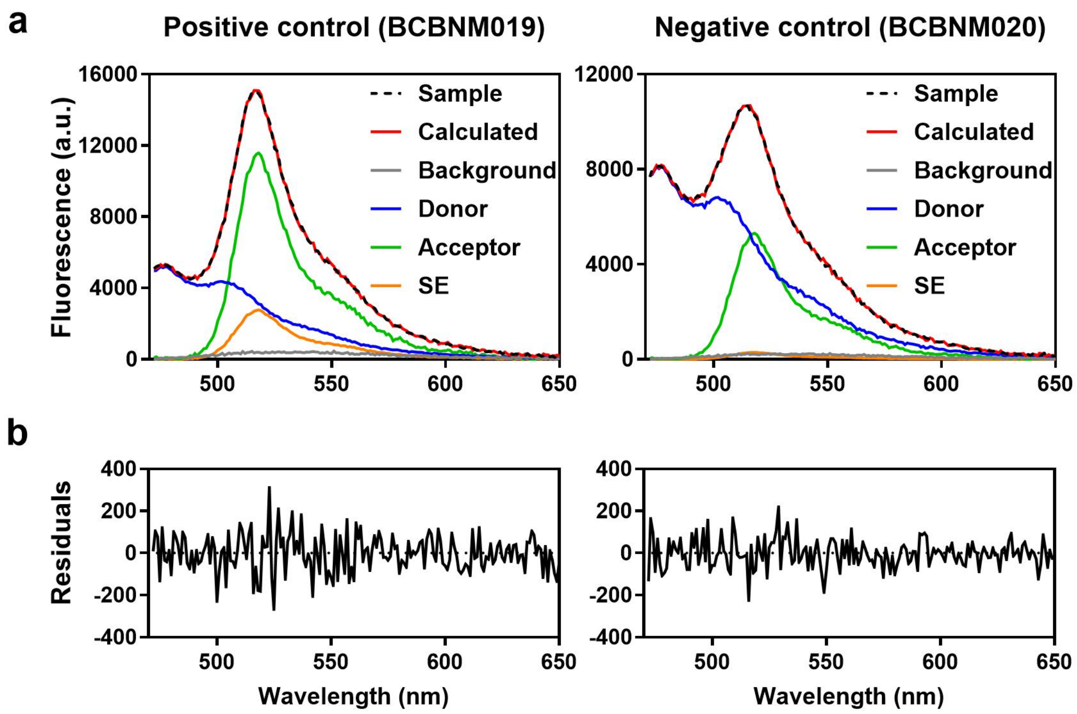

Figure S9 – Representative spectral unmixing data of fluorescence spectra measured for the Zip-Zip interaction control strains.

**a**) Unmixing of the measured acceptor spectra into their individual spectral components to determine acceptor based FRET (**see Methods**) performed as described in Meiresonne *et al* [7]. Reference spectra were obtained from strains BCBNM016 (donor), BCBNM017 (acceptor) and JE2 (background). The black dashed lines show the measured spectrum which is dissected into a spectrum for the background of JE2 cells (gray), sfTq2$^{ox}$ (blue), mNG (green) and the sensitized emission (SE, orange). The sum of the unmixed spectra is represented as a red line (calculated). The calculated FRET efficiencies were 16.2 ± 2.0 % (n = 2) for the positive control (BCBNM019) and 2.6 ± 0.0 % (n = 4) for the negative control (BCBNM020). **b**) Residual signal that could not be accounted for by unmixing is plotted (measured signal – calculated signal) and served as a measure of quality.

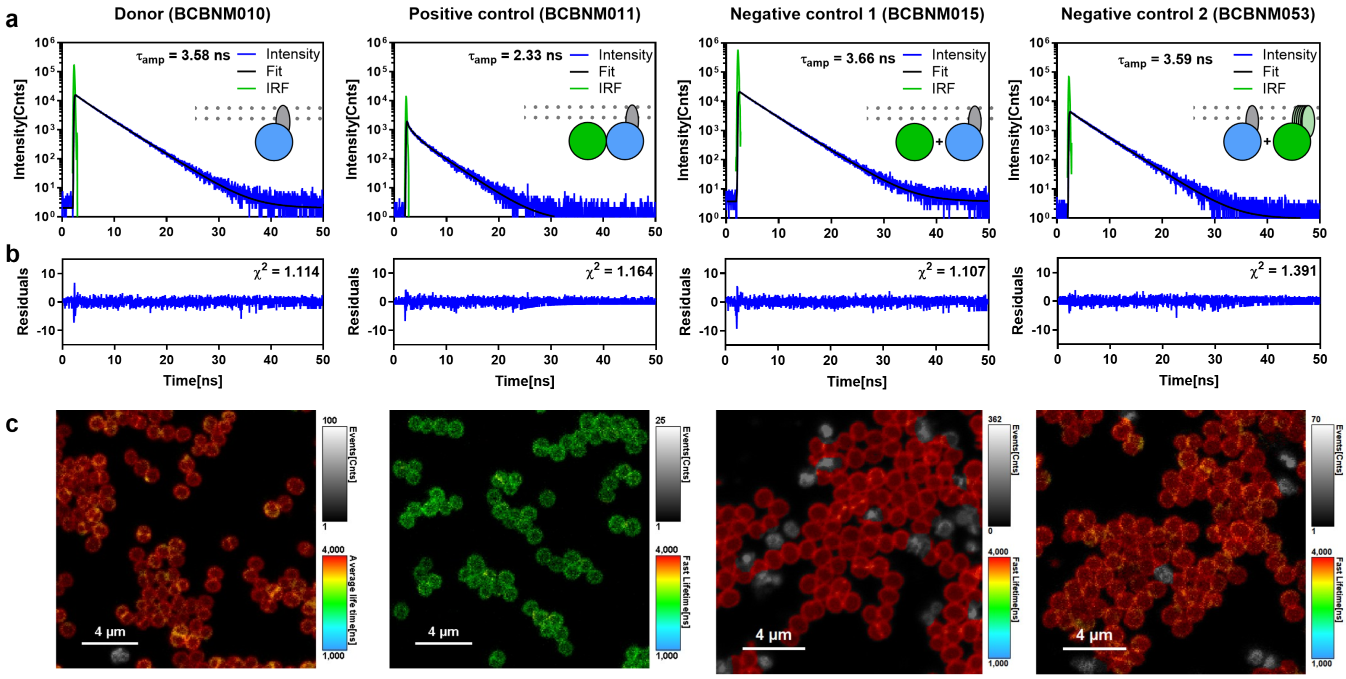

#### Figure S10 – Representative fluorescence decay curves for control strains expressing membrane-anchored fluorescent proteins.

**a**) Representative fluorescence decay curves for the indicated JE2 derived strains expressing sfTq2$^{ox}$-TM (Donor), mNG-sfTq2$^{ox}$-TM (Positive control), mNG + sfTq2$^{ox}$-TM (Negative control 1) and sfTq2$^{ox}$-TM + Lrp-mNG (Negative control 2). Cartoons represent fluorescent proteins produced in each strain. $\tau_{amp}$ = amplitude weighted average fluorescence lifetime, IRF = Instrument Response Function. **b**) Residuals corresponding to the fit of the fluorescence decay curves presented in a. **c**) FastFLIM (**see Methods**) images corresponding to the fluorescence decay curves shown in a.

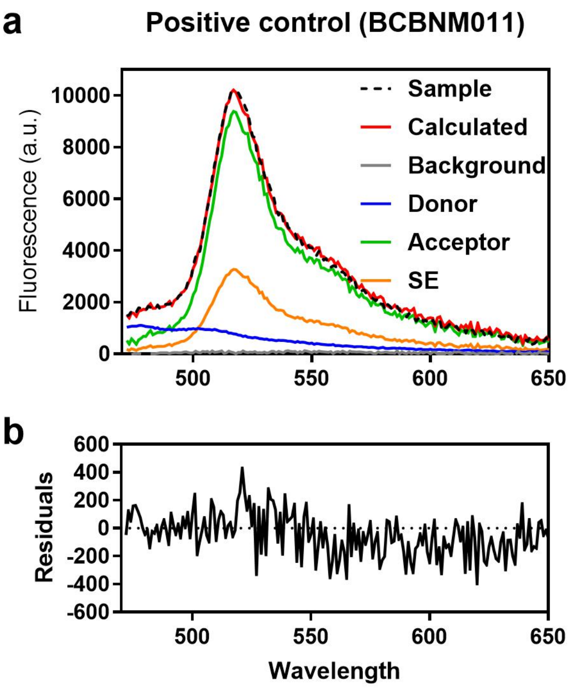

Figure S11 –Representative spectral unmixing of fluorescence spectrum measured for the positive control strain producing membrane anchored TM-mNG-sfTq2$^{ox}$.

**a**) Unmixing of the measured acceptor spectra into their individual spectral components to determine acceptor based FRET (**see Methods**) performed as described in Meiresonne *et al* [7]. Reference spectra were obtained from strains BCBNM010 (donor), BCBNM009 (acceptor) and JE2 (background). The black dashed line shows the measured spectrum which is dissected into a spectrum for the background of JE2 cells (gray), sfTq2$^{ox}$ (blue), mNG (green) and the sensitized emission (SE, orange). The sum of the unmixed spectra is represented as a red line (calculated). The calculated FRET efficiency for the positive control (BCBNM003) was 30.0 ± 2.8 % (n = 9). **b**) Residual signal that could not be accounted for by unmixing is plotted (measured signal – calculated signal) and served as a measure of quality.

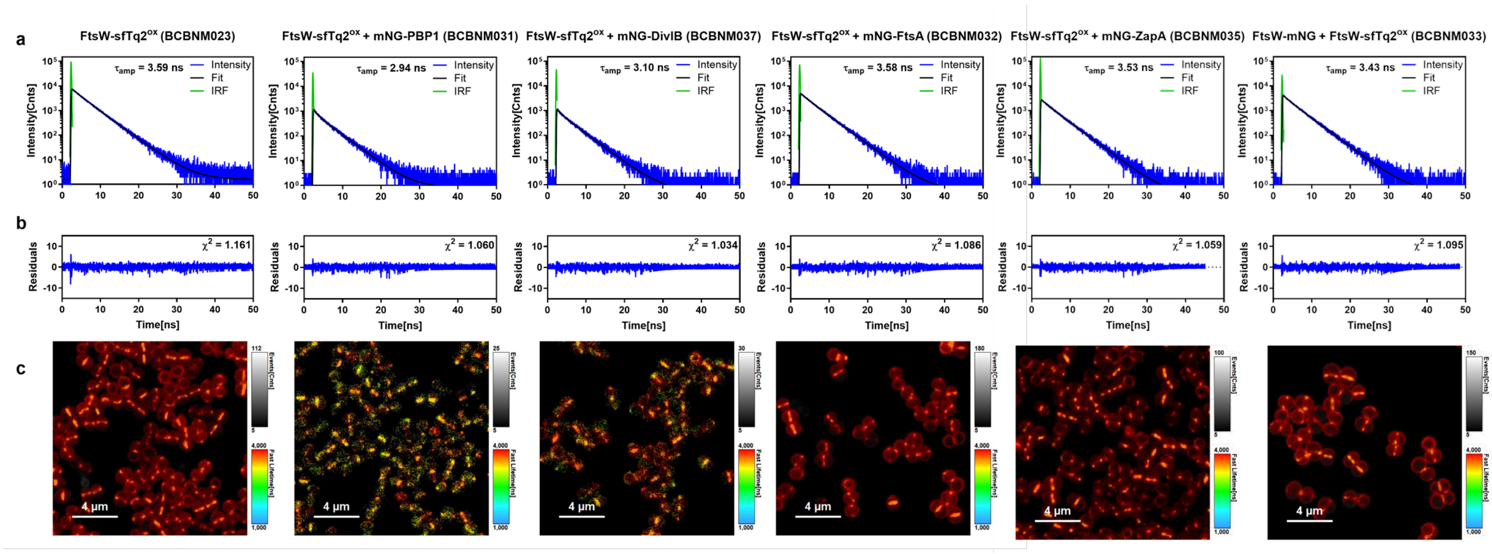

### Figure S12 – Representative decay curves for the strains to assess FtsW-sfTq2$^{ox}$ protein-protein interactions

**a**) Representative decay curves for the indicated strains producing FtsW-sfTq2$^{ox}$ or FtsW-sfTq2$^{ox}$ and a mNG derivative of cell division proteins of interest PBP1, DivIB, FtsA, ZapA or FtsW itself. $\tau_{amp}$ = amplitude weighted average fluorescence lifetime, IRF = Instrument Response Function. **b**) Residuals corresponding to the fit of the fluorescence decay curves presented in a. **c**) FastFLIM (**see Methods**) images corresponding to the fluorescence decay curves shown in a.

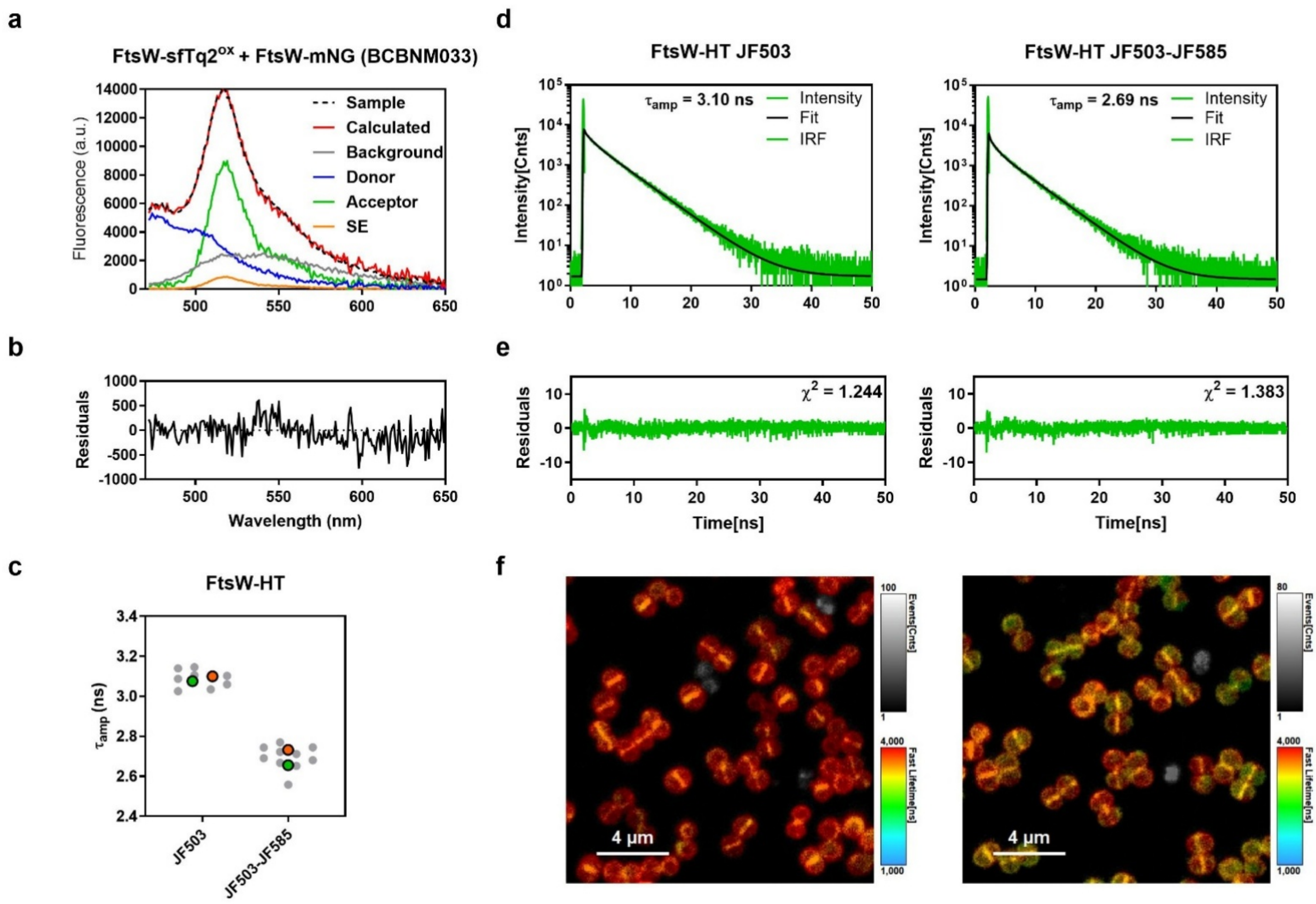

#### Figure S13 – Supporting evidence for a FtsW-FtsW interaction.

**a**) Unmixing of the measured acceptor spectrum for strain BCBNM033, producing FtsW-sfTq2$^{ox}$ and FtsW-mNG, into its individual spectral components to determine acceptor-based FRET (**see Methods**) performed as described in Meiresonne *et al* [7]. Reference spectra were obtained from strains BCBNM023 (FtsW-sfTq2$^{ox}$, donor), BCBNM022 (FtsW-mNG, acceptor) and JE2 (background). The black dashed line shows the measured spectrum which is dissected into a spectrum for the background of JE2 cells (gray), sfTq2$^{ox}$ (blue), mNG (green) and the sensitized emission (SE, orange). The sum of the unmixed spectra is represented as a red line (calculated). The E*f*A value was 7.1 ± 0.9 % (n = 5) for the FtsW-FtsW interaction in strain BCBNM033. **b**) Residual signal that could not be accounted for by unmixing is plotted (measured signal – calculated signal) and serves as a measure of quality. **c**) Superplot of amplitude weighted average fluorescence lifetimes measured in strain BCBSS187, producing FtsW-HT, labelled with JF503 or a 1:1 mix of JF503 (donor) and JF585 (acceptor). Each grey dot represents the fitted lifetime of a single FLIM image. The coloured dots represent the average of the FLIM images in an individual experiment. **d**) Representative fluorescence decay curves of the data presented in c. **e**) Residuals of the fitted decay curves from d. **f**) FastFLIM (**see Methods**) images of strain BCBSS187 labelled with JF503 (left) or a 1:1 mix of JF503 and JF585 (right).

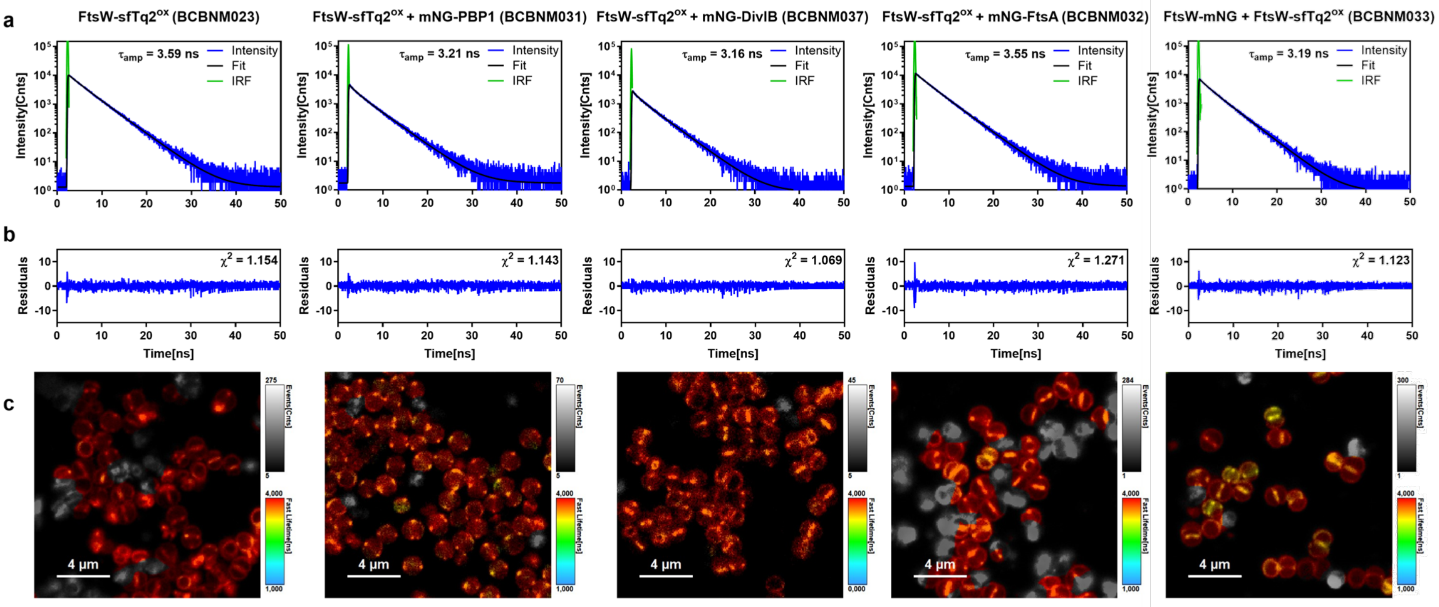

#### Figure S14 – Representative decay curves for the strains to assess FtsW-sfTq2$^{ox}$ protein-protein interactions in the presence of imipenem

**a)** Representative decay curves for the indicated strains producing FtsW-sfTq2$^{ox}$ or FtsW-sfTq2$^{ox}$ and a mNG derivative of cell division proteins of interest PBP1, DivIB, FtsA treated with 1 µg / ml imipenem. $\tau_{amp}$ = amplitude weighted average fluorescence lifetime, IRF = Instrument Response Function. **b**) Residuals corresponding to the fit of the fluorescence decay curves presented in a. **c**) FastFLIM (**see Methods**) images corresponding to the fluorescence decay curves shown in a. Signals from the greyed-out regions were excluded from fitting analysis.

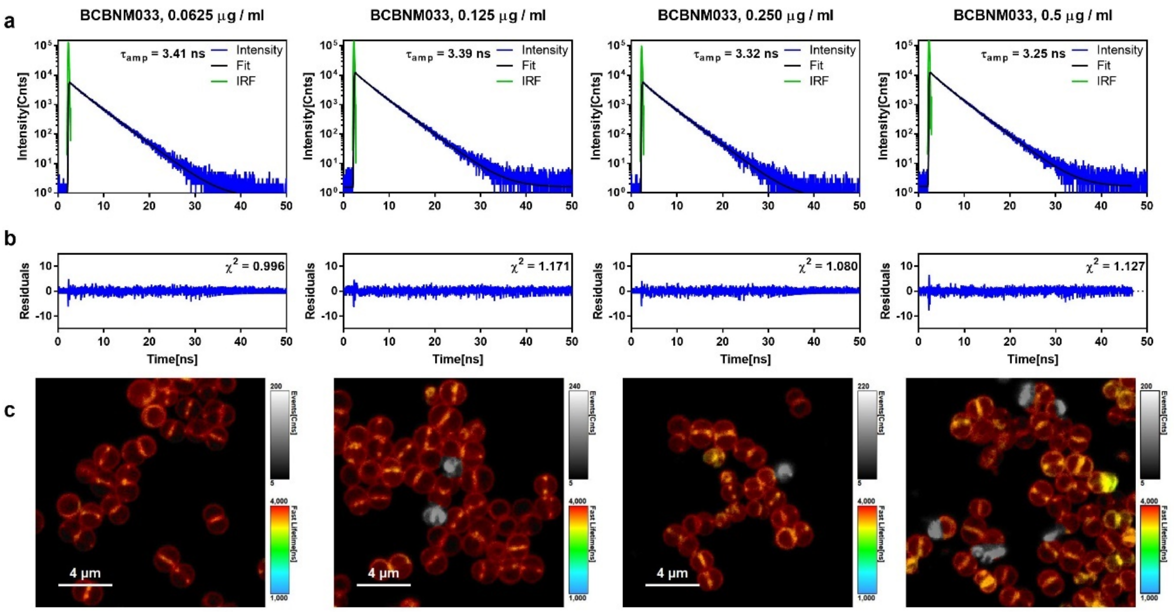

#### Figure S15 – Representative decay curves for strain used to assess the FtsW-sfTq2$^{ox}$ – FtsW-mNG interaction with different concentrations of imipenem.

**a)** Representative decay curves for strain BCBNM033 producing FtsW-mNG and FtsW-sfTq2$^{ox}$, treated with 0.0625, 0.125, 0.250 and 0.500 µg / ml imipenem. Data for 0 µg / ml imipenem is given in panel a of Figure S14. $\tau_{amp}$ = amplitude weighted average fluorescence lifetime, IRF = Instrument Response Function. **b**) Residuals corresponding to the fit of the fluorescence decay curves presented in a. **c**) FastFLIM (**see Methods**) images corresponding to the fluorescence decay curves shown in a. Signals from the greyed-out regions were excluded from fitting analysis.

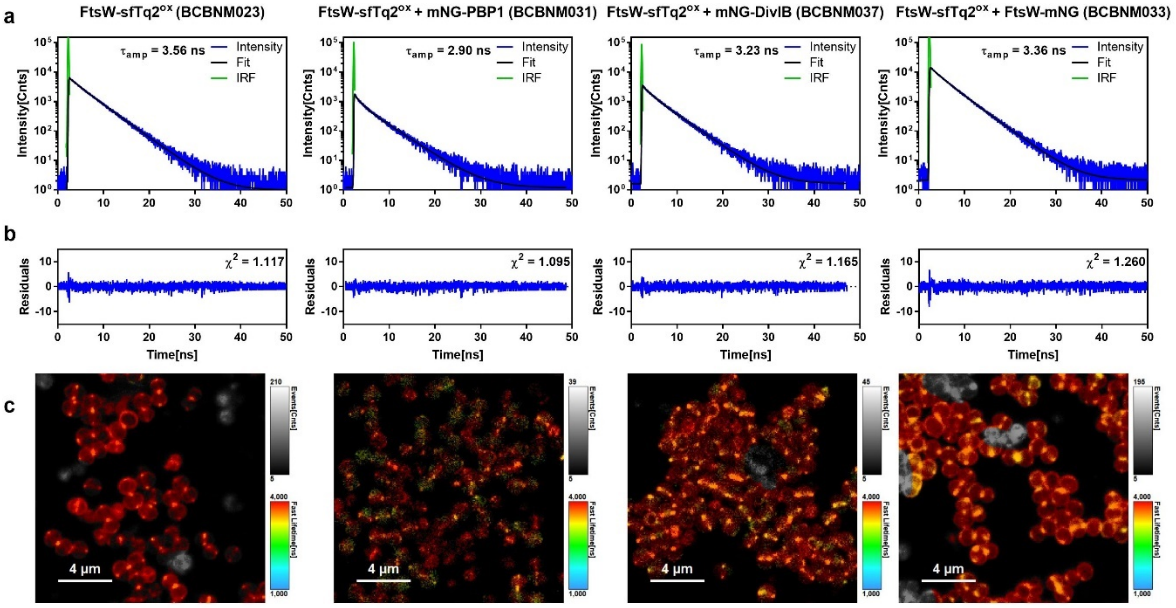

#### Figure S16 – Representative decay curves for the strains to assess FtsW-sfTq2$^{ox}$ protein-protein interactions in the presence of DMPI

**a)** Representative decay curves for the indicated strains producing FtsW-sfTq2$^{ox}$ or FtsW-sfTq2$^{ox}$ and a mNG derivative of cell division proteins PBP1, DivIB or FtsW itself, treated with 8 µg / ml DMPI. $\tau_{amp}$ = amplitude weighted average fluorescence lifetime, IRF = Instrument Response Function. **b**) Residuals corresponding to the fit of the fluorescence decay curves presented in a. **c**) FastFLIM (**see Methods**) images corresponding to the fluorescence decay curves shown in a. Signals from the greyed-out regions were excluded from fitting analysis.

### Table S1 – Strains used in this study

| Name | Plasmid used | Genotype | Description | Ref |
|---|---|---|---|---|
| ***Escherichia coli*** | | | | |
| DC10B | - | Δ*dcm* in DH10B background; Dam methylation only | Cloning strain | [8] |
| DH5α | - | F$^-$, *end*A1, hsdR17 (r$^-$, m$^+$), *sup*E44, *thi*-1, λ-, *rec*A1, *gyr*A96, *rel*A1, ϕ80d*lac*ZΔM15 | Cloning strain | [9] |
| BL21 (DE3) | - | B F$^-$ *dcm ompT hsdS* ($r_B^-m_B^-$) *gal* λ (DE3) | Protein purification | Stratagene |
| ***Staphylococcus aureus*** | | | | |
| RN4220 | - | Restriction-negative derivative of NCTC8325-4 | | [10][11] |
| JE2 | - | CA-MRSA; | WT; Background reference | [12] |
| COL | - | HA-MRSA | WT | [13] |
| **Reference and control strains for interaction of cytosolic proteins** | | | | |
| BCBNM001 | pBCBNM001 | JE2 *spa::Pxyl-mNG* | JE2 producing mNG; acceptor reference | This study |
| BCBNM002 | pBCBNM002 | JE2 *spa::Pxyl-sfTq2$^{ox}$* | JE2 producing sfTq2$^{ox}$; donor reference | This study |
| BCBNM003 | pBCBNM003 | JE2 *spa::Pxyl-mNG-sfTq2$^{ox}$* | JE2 producing mNG-sfTq2$^{ox}$; positive control | This study |
| BCBNM005 | pBCBNM005 | JE2 *spa::Pxyl-sfTq2$^{ox}$ + mNG* | JE2 producing sfTq2$^{ox}$ and mNG; negative control | This study |
| **Reference and control strains for Zip-Zip interaction** | | | | |
| BCBNM016 | pBCBNM016 | JE2 *spa::Pxyl-sfTq2$^{ox}$-zip$^{25}$* | JE2 producing sfTq2$^{ox}$-Zip$^{25}$; donor reference | This study |
| BCBNM017 | pBCBNM017 | JE2 *spa::Pxyl-mNG-zip$^{18}$* | JE2 producing mNG-Zip$^{18}$; acceptor reference | This study |
| BCBNM019 | pBCBNM019 | JE2 *spa::Pxyl-mNG-zip$^{18}$ + sfTq2$^{ox}$-zip$^{25}$* | JE2 producing mNG-Zip$^{18}$ and sfTq2$^{ox}$-Zip$^{25}$; positive control | This study |
| BCBNM020 | pBCBNM020 | JE2 *spa::Pxyl-sfTq2$^{ox}$-zip$^{25}$ + mNG* | JE2 producing sfTq2$^{ox}$-Zip$^{25}$ and mNG; negative control | This study |
| **Reference and control strains for interaction of membrane associated proteins** | | | | |
| BCBNM009 | pBCBNM009 | JE2 *spa::Pxyl-3tetO-mNG-TM* | JE2 producing mNG-TM; acceptor reference | This study |
| BCBNM010 | pBCBNM010 | JE2 *spa::Pxyl-3tetO-sfTq2$^{ox}$-TM* | JE2 producing sfTq2$^{ox}$-TM; donor reference | This study |
| BCBNM011 | pBCBNM011 | JE2 *spa::Pxyl-3tetO-mNG-sfTq2$^{ox}$-TM* | JE2 producing mNG-sfTq2$^{ox}$-TM; positive control | This study |
| BCBNM015 | pBCBNM015 | JE2 *spa::Pxyl-mNG + sfTq2$^{ox}$-TM* | JE2 producing mNG and sfTq2$^{ox}$-TM; negative control 1 | This study |
| BCBNM053 | pBCBSS206 | JE2 *spa::Pxyl-sfTq2$^{ox}$-TM + SAUSA300_2100::SAUSA300_2100-mNG* | JE2 producing sfTq2$^{ox}$-TM and *SAUSA300_2100*-mNG; negative control 2 | This study |
| **Strains for assessing FtsW interactions** | | | | |
| BCBNM023 | pBCBNM023 | JE2 *spa::Pxyl-ftsW-sfTq2$^{ox}$* | JE2 producing FtsW-sfTq2$^{ox}$; donor reference | This study |
| BCBNM022 | pBCBNM022 | JE2 *spa::Pxyl-ftsW-mNG* | JE2 producing FtsW-mNG; acceptor reference | This study |
| BCBNM031 | pBCBNM031 | JE2 *spa::Pxyl-mNG$^{ecySA}$-pbp1 + ftsW-sfTq2$^{ox}$* | JE2 producing mNG-PBP1 and FtsW-sfTq2$^{ox}$; Strain to assess the interaction between FtsW and PBP1 | This study |
| BCBNM037 | pBCBNM037 | JE2 *spa::Pxyl-mNG$^{ecySA}$-divIB + ftsW-sfTq2$^{ox}$* | JE2 producing mNG-DivIB and FtsW-sfTq2$^{ox}$; Strain to assess the interaction between FtsW and DivIB | This study |
| BCBNM033 | pBCBNM033 | JE2 *spa::Pxyl-ftsW-mNG + ftsW-sfTq2$^{ox}$* | JE2 producing FtsW-mNG and FtsW-sfTq2$^{ox}$; Strain to assess the FtsW self-interaction | This study |
| BCBNM032 | pBCBNM032 | JE2 *spa::Pxyl-ftsW-sfTq2$^{ox}$ + mNG$^{ecySA}$-ftsA* | JE2 producing mNG-FtsA and FtsW-sfTq2$^{ox}$; Strain to assess the interaction between FtsW and FtsA | This study |
| BCBNM035 | pBCBNM035 | JE2 *spa::PXyl-mNG$^{ecySA}$-zapA + FtsW-sfTq2$^{ox}$* | JE2 producing mNG-ZapA and FtsW-sfTq2$^{ox}$; Strain to assess the interaction between FtsW and ZapA | This study |
| BCBSS187 | pBCBSS074 | JE2 *spa::ftsW-halo* | JE2 producing FtsW-halo; Strain to assess FtsW self-interaction and donor reference (depending on the labelling) | This study |

### Table S2 – Plasmids used and cloning strategies

| Short name | Long name | Information | Ref. | Vector backbone | RE1* | RE2* | Insert | Template | Primers | RE3* | RE4* |
|---|---|---|---|---|---|---|---|---|---|---|---|
| **Miscellaneous and cloning plasmids** | | | | | | | | | | | |
| **pBCBNM000** | pBCB-Pxyl-st3-mNG-2GGS-MurJ-2GGS-sfTq2$^{ox}$ | Made by Gibson assembly. Used as a cloning vector by replacing MurJ with insert of interest | This study | pBCB43 | EagI | | mNG / MurJ / sfTq2$^{ox}$ | pBCBSMC018 / JE2 chromosome / pBCBSMC017 | 11647 + 11648 / 11649 + 11650 / 11651 + 11652 | | |
| **pBCB13** | - | pMAD derivative with up- and downstream regions of the *spa* locus and P*spac-lacI*, | [14] | | | | | | | | |
| **pMadΔspa** | - | pMAD derivative with the up- and downstream regions of spa to delete spa gene | [15] | | | | | | | | |
| **pBCB43** | - | pBCB13 derivative lacking P*spac* and *lacI*, containing *tetR*, Pxyl *tetO*, and a downstream located *tetO* site, *lacZ* | [16] | | | | | | | | |
| **pBCB36** | pBCB13-Pxyl-sfGFP | pBCB13 derivative with Pxyl promoter N25, without the promoter for the repressor making the product highly expressed (constitutive), Used to amplify N25-sfGFP | Nathalie Reichmann | | | | | | | | |
| **pCNX** | - | Replicative vector containing the cadmium-inducible P*cad* promoter, Used as a cloning vector | [17] | | | | | | | | |
| **pET28a-mTq2** | - | Commercially ordered. Used for mTq2 protein purification | This study | | | | | | | | |
| **pET28a-sfTq2$^{ox}$** | - | Commercially ordered. Used for sfTq2$^{ox}$ protein purification | This study | | | | | | | | |
| **pCNX-mCh-sGFP-TM** | - | Used to amplify TM | [6] | | | | | | | | |
| **pBCB33-Neon-Z** | pBCB13-Pcad-mNeonGreen-15aaBS-FtsZ | Used to amplify mNG | [18] | | | | | | | | |
| **pBCBSS135** | pBCB13-3XFLAG-mNeonGreen | Used to amplify mNG | [15] | | | | | | | | |
| **pBCBHV004** | pMutin-Spo0J-yfp | Used as a cloning vector by replacing Spo0J by insert of interest | [19] | | | | | | | | |
| **pJ201mNeonGreen** | - | Commercially ordered. Used to amplify mNG | Raquel Pereira | | | | | | | | |
| **pUT18c** | - | Used to amplify zip$^{18}$ | [20] | | | | | | | | |
| **pKT25** | - | Used to amplify zip$^{25}$ | [20] | | | | | | | | |
| **pBCBMF019** | pCNX-FtsW-sfTq2$^{ox}$ | Made by Gibson assembly. Used to amplify FtsW-sfTq2$^{ox}$ | This study | pCNX | SmaI | - | ftsW / sfTq2$^{ox}$ | COL chromosome / pBCBSMC017 | 7395 + 7934 / 7935 + 7938 | - | - |
| **pBCBMF013** | pCNX-FtsW-mNG | Made by Gibson assembly. Used as a cloning template to amplify mNG or FtsW-mNG | This study | pCNX | SmaI | - | ftsW / mNG | COL chromosome / pJ201mNeonGreen | 7395 + 7390 / 7391 + 7396 | - | - |
| **pBCBMF004** | pBCB13-FtsA-mNG | Made by Gibson assembly. Used to amplify mNG-FtsA | | pBCB13 | XhoI | - | ftsA / mNG | COL chromosome / pJ201mNeonGreen | 6819 +6820 / 6817 + 6818 | - | - |
| **pBCBMF036** | pCNX-mNG-FtsA | Used to amplify mNG$^{ecySA}$-FtsA | This study | pCNX | SalI | KpnI | mNG-ftsA | pBCBMF004 | 8702 + 8709 | SalI | KpnI |
| **pBCBMF003** | pBCB13-mNG-ZapA | Made by Gibson assembly. Used to amplify mNG-ZapA | | pBCB13 | XhoI | - | zapA / mNG | COL chromosome / pJ201mNeonGreen | 6815 + 6816 / 6817 + 6818 | - | - |
| **pBCBMF035** | pCNX-mNG-ZapA | Used to amplify mNG$^{ecySA}$-ZapA | This study | pCNX | SalI | EcoRI | mNG-ZapA | pBCBMF003 | 8702 + 8703 | SalI | EcoRI |
| **pBCBSMC018** | pCNX-mNG-TM | Made by Gibson assembly. Used to amplify mNG or mNG-TM | This study | pCNX | SmaI | - | mNG / TM | pBCB33-Neon-Z / pCNX-mCh-sGFP-TM | 7068 + 7071 / 6400 + 7075 | - | - |
| **pBCBSMC016** | pCNX-mNG-sfTq2$^{ox}$-TM | Made by Gibson assembly. Used to amplify sfTq2$^{ox}$-TM or mNG-sfTq2$^{ox}$-TM | This study | pCNX | SmaI | - | mNG / sfTq2$^{ox}$ / TM | pBCB33-Neon-Z / sfTq2$^{ox}$** /pCNX-mCh-sGFP-TM | 7070 + 7077 / 7068 + 7071 / 6400 + 7075 | - | - |
| **pBCBNM007** | pMAD-Spa-Pxyl-sfGFP | Used as a cloning vector | This study | pMadΔspa | EcoRI | NheI | N25-sfGFP | pBCB36 | 8435 + 8436 | EcoRI | NheI |

| | | | | | | | | | | | |
|---|---|---|---|---|---|---|---|---|---|---|---|
| **pBCBNM024** | pBCB-Pxyl 3tetO-st7-mNG$^{ecySA}$-FtsA | Used to amplify mNG$^{ecySA}$-FtsA | This study | pBCBNM000 | TspMI | NheI | mNG$^{ecySA}$-FtsA | pBCBMF036 | 9640 + 9639 | TspMI | NheI |
| **pBCBNM025** | pBCB-Pxyl 3tetO-st7-mNG$^{ecySA}$-ZapA | Used to amplify mNG$^{ecySA}$-ZapA | This study | pBCBNM000 | TspMI | NheI | mNGe$^{cySA}$-ZapA | pBCBMF035 | 9641 + 9640 | TspMI | NheI |
| **pBCBNM027** | pBCB-Pxyl 3tetO-st7-mNG$^{ecySA}$-PBP1 | Insert by overlap extension PCR. Used as to amplify mNG$^{ecySA}$-PBP1, | This study | pBCBNM000 | TspMI | NheI | mNG$^{ecySA}$-PBP1 | pBCBMF013 / JE2 chromosome | 9640 + 9646 / 9645 + 9647 | TspMI | XbaI |
| **pBCBNM029** | pBCB-Pxyl 3tetO-st7-mNG$^{ecySA}$-DivIB | Insert by overlap extension PCR. Used to amplify mNG$^{ecySA}$-DivIB, | This study | pBCBNM000 | TspMI | NheI | mNG$^{ecySA}$-DivIB | pBCBMF013 / JE2 chromosome | 9640 + 9652 / 9651 + 9653 | TspMI | NheI |
| **Cytoplasmic Expression constructs** | | | | | | | | | | | |
| **pBCBNM001** | pMAD-spa-Pxyl-mNG | Used to create strain BCBNM001. Used to amplify mNG | This study | pBCBNM007 | SmaI | NheI | mNG | pBCBSMC018 | 8439 + 8440 | SmaI | NheI |
| **pBCBNM002** | pMAD-spa-Pxyl-sfTq2$^{ox}$ | Used to create strain BCBNM002. Used to amplify sfTq2$^{ox}$ | This study | pBCBNM007 | SmaI | NheI | sfTq2$^{ox}$ | pBCBSMC017 | 8441 + 8442 | SmaI | NheI |
| **pBCBNM003** | pMAD-spa-Pxyl-mNG-sfTq2$^{ox}$ | Used to create strain BCBNM003 | This study | pBCBNM007 | SmaI | NheI | mNG-sfTq2$^{ox}$ | pBCBSMC016 | 8439 + 8442 | SmaI | NheI |
| **pBCBNM005** | pMAD-spa-pXyl-sfTq2$^{ox}$_mNG | Used to create strain BCBNM005 | This study | pBCBNM002 | NheI | - | mNG | pBCBNM001 | 11653 + 8443 | NheI | - |
| **Zip-Zip Expression constructs** | | | | | | | | | | | |
| **pBCBNM016** | pMAD-spa-Pxyl-sfTq2$^{ox}$-Zip$^{25}$ | Insert by overlap extension PCR. Used to create strain BCBNM016. Used to amplify Zip$^{25}$ | This study | pBCBNM007 | TspMI | NheI | sfTq2$^{ox}$-Zip$^{25}$ | pBCBNM002 / pKT25 | 8441 + 9252 / 9251 + 9253 | TspMI | NheI |
| **pBCBNM017** | pMAD-spa-Pxyl-mNG-Zip$^{18}$ | Insert by overlap extension PCR. Used to create strain BCBNM017. Used to amplify Zip$^{18}$ | This study | pBCBNM007 | TspMI | NheI | mNG-Zip$^{18}$ | pBCBNM001 / pUT18c | 9255 +8439 / 9254 + 9256 | TspMI | NheI |
| **pBCBNM019** | pMAD-spa-Pxyl-mNG-Zip18_sfTq2$^{ox}$-Zip25 | Used to create strain BCBNM019 | This study | pBCBNM017 | NheI | - | sfTq2$^{ox}$-Zip$^{25}$ | pBCBNM016 | 8443 + 9253 | NheI | - |
| **pBCBNM020** | pMAD-spa-Pxyl-sfTq2$^{ox}$-Zip25_mNG-cyto | Used to create strain BCBNM020 | This study | pBCBNM016 | TspMI | NheI | mNG | pBCBNM017 | 8440 + 8443 | | |
| **TM Expression constructs** | | | | | | | | | | | |
| **pBCBSMC017** | pCNX-sfTq2$^{ox}$-TM | Made by Gibson assembly | This study | pCNX | SmaI | - | sfTq2$^{ox}$ / TM | sfTq2$^{ox**}$ / pCNX-mCh-sGFP-TM | 7070 + 7077 / 6400 + 7075 | - | - |
| **pBCBNM009** | pBCB-Pxyl-3tetO-st7-mNG-TM | Used to create strain BCBNM009 | This study | pBCBNM000 | TspMI | NheI | mNG-TM | pBCBSMC018 | 8867 + 8863 | TspMI | NheI |
| **pBCBNM010** | pBCB-Pxyl-3tetO-st7-sfTq2$^{ox}$-TM | Used to create strain BCBNM010. Used to amplify sfTq2$^{ox}$-TM | This study | pBCBNM000 | TspMI | NheI | sfTq2$^{ox}$-TM | pBCBSMC016 | 8865 + 8863 | TspMI | NheI |
| **pBCBNM011** | pBCB-Pxyl-3tetO-st7-mNG-sfTq2$^{ox}$-TM | Used to create strain BCBNM011 | This study | pBCBNM000 | TspMI | NheI | mNG-sfTq2$^{ox}$-TM | pBCBSMC016 | 8867 + 8863 | TspMI | NheI |
| **pBCBNM015** | pBCB-Pxyl-mNGcyto-st7-sfTq2$^{ox}$-TM | Used to create strain BCBNM015 | This study | pBCBNM001 | NheI | - | sfTq2$^{ox}$-TM | pBCBNM010 | 8440 + 8439 | NheI | - |
| **pBCBSS206** | pMutin-SAUSA300_2100-mNG | Used to create strain BCBNM053 | This study | pBCBHV004 | SpeI | HindIII | Lrp-mNG | JE2 chromosome / pBCBSS135 | 8296 + 8297 / 7359 + 8295 | HindIII /SalI | SalI/ NheI |
| **FtsW interactome Expression constructs** | | | | | | | | | | | |
| **pBCBNM023** | pBCB-pXyl-3tetO-st7-FtsW-sfTq2$^{ox}$ | Used to create strain BCBNM023. Used to amplify FtsW-sfTq2$^{ox}$ | This study | pBCBNM000 | TspMI | NheI | FtsW-sfTq2$^{ox}$ | pBCBMF019 | 9638 + 8442 | TspMI | NheI |
| **pBCBNM022** | pBCB-pXyl-3tetO-st7-FtsW-mNG | Used to create strain BCBNM022 | This study | pBCBNM000 | TspMI | NheI | FtsW-mNG | pBCBMF013 | 9638 + 8442 | TspMI | NheI |
| **pBCBNM031** | pBCB-pXyl-3tetO-st7-mNG$^{ecySA}$-PBP1_FtsW-sfTq2$^{ox}$ | Used to create strain BCBNM031 | This study | pBCBNM027 | EagI | - | FtsW-sfTq2$^{ox}$ | pBCBNM023 | 9680 + 9679 | EagI | - |
| **pBCBNM037** | pBCB pXyl-3tetO-st7-mNG$^{ecySA}$-DivIB_FtsW-sfTq2$^{ox}$ | Used to create strain BCBNM037 | This study | pBCBNM029 | NheI | - | FtsW-sfTq2$^{ox}$ | pBCBNM023 | 9729 + 8442 | NheI | - |
| **pBCBNM033** | pBCB-pXyl-3tetO-st7-FtsW-mNG_FtsW-sfTq2$^{ox}$ | Used to create strain BCBNM033 | This study | pBCBNM022 | NheI | - | FtsW-sfTq2$^{ox}$ | pBCBNM023 | 9729 + 8442 | NheI | - |
| **pBCBNM032** | pBCB-pXyl-3tetO-st7-FtsW-sfTq2$^{ox}$_mNG$^{ecySA}$-FtsA | Used to create strain BCBNM032 | This study | pBCBNM023 | NheI | - | mNG$^{ecySA}$-FtsA | pBCBNM024 | 9639 + 9681 | NheI | - |
| **pBCBNM035** | pBCB-pXyl-3tetO-st7-mNG$^{ecySA}$-ZapA_FtsW-sfTq2$^{ox}$ | Used to create strain BCBNM035 | This study | pBCBNM025 | NheI | - | FtsW-sfTq2$^{ox}$ | pBCBNM023 | 9729 +8442 | NheI | - |
| **pBCBSS074** | pBCB43-ftsWht | Used to create strain BCBSS187 | This study | pBCB43 | SmaI | EagI | FtsW-HT | JE2 EzrA-sGFP FtsW-HT chromosome [21] | 7034 + 7141 | SmaI | EagI |

* Restriction enzymes RE1 and RE2 were used to digest the vector backbone and RE3 and RE4 were used to digest inserts for cloning. When cloning was done by Gibson assembly, as indicated, RE1 was used to linearize the vector construct. **sfTq2$^{ox}$ synthetic dsDNA ordered from IDT. mNG$^{ecySA}$ is a mNG derivative (mNG$^{\Delta 1\text{-}9}$) optimised for improved expression and N-terminal FP fusion production [22].

#### Table S3 – Primers used

| Short Name | Long Name | Oligo sequence 5' → 3' |
|---|---|---|
| 6400 | GGSGGGGS-TM2_fwd | GGAGGCGGTTCTGGCGGAGGTGGCTCTACGGAAAACAAAGGATCTTCTCAG |
| 6815 | mNeon_link_zapA_fwd | TCCTGCGGAGCCTCTATGACACAGTTTAAAAACAAGG |
| 6816 | zapA_pBCB13_rev | GCGGCGCCTCGGAGCGACTTATTGCTCACGCTGCTGC |
| 6817 | pBCB13_rbs_mNeon_fwd | GGGTTTCTAGAGCCTAGCAGGAGGACTCGGATGGTATCAAAAGG |
| 6818 | mNeon_link_2mut_rev | AGAGGCTCCGCAGGATTTGTATAACTCATCCATGC |
| 6819 | mNeon_link_ftsA_fwd | TCCTGCGGAGCCTCTATGGAAGAACATTACTACG |
| 6820 | ftsA_pBCB13_rev | GCGGCGCCTCGGAGCGACTCATTCAAATAGAGATTTCATTAG |
| 7034 | C-halo-tag_rev2 | ATATCTCGAGCGGCCGTTAACCACTGATTTCTAAAGTAGATAACCATC |
| 7068 | pCNX-RBS-mNG | GTCGACTCTAGAGGATCCCCAATATCTAAGGAGGTAATATAATGGTATCAAAAGGTGAAG |
| 7070 | pCNX-RBSsftq2$^{ox}$ | GTCGACTCTAGAGGATCCCCAATATCTAAGGAGGTAATATAATGAGTAAAGGCGAAGAG |
| 7071 | mNG_linker_sfTq2$^{ox}$_Rev | CTTTACTACCCAATTCTTTGTATAACTCATCC |
| 7072 | mNG_linker_mtq2_fwd | CAAAGAATTGGGTAGTAAAGGCGAAGAG |
| 7075 | TM-pCNX | CTGAATTCGAGCTCGGTACCCTTAAGCAAACAATAAGATACC |
| 7077 | mNG/sfTq2oxlinker rev | CACCTCCGCCAGAACCGCCTCCACCTTTGTATAACTCATCCATGC |
| 7141 | COLftsW_st7_SmaI_fwd | ATATCCCGGGAAAAAATAAGGAGGAAAAAAAATGAAGAATTTTAGAAGTATTTTACG |
| 7359 | MCS_SCGAS_mNG_+4_fwd | ATATCCCGGGCGGCCGAGTACTGTCGACTCCTGCGGCGCCTCCGTATCAAAAGGTGAAGAAGATAATATGGC |
| 7390 | P6_ftsW-mNG_pBCB13 | CTTCACCTTTTGATACCATAGAACCTCCTCCACCAGAACCTCCTCCACC |
| 7391 | P7_ftsW-mNG_pBCB13 | TCTGGTGGAGGAGGTTCTATGGTATCAAAAGGTGAAGAAG |
| 7394 | P10_sfTq-PBP1_pCNX | AATTCGAGCTCGGTACCCTTAGTCCGACTTATCCTTGTC |
| 7395 | P11_ftsW-mNG_pCNX | CGACTCTAGAGGATCCCCAGGAGGAAATGAAATGAAGAATTTTAGAAG |
| 7396 | P12_ftsW-mNG_pCNX | AATTCGAGCTCGGTACCCTTATTTGTATAACTCATCCATGC |
| 7935 | P13_ftsW-sfTq_pCNX | GGAGGTTCTATGAGTAAAGGCGAAGAG |
| 7938 | P14_ftsW-sfTq_pCNX | AATTCGAGCTCGGTACCCTTATTTGTATAACTCATCCATGC |
| 8295 | mNG_Nhe_rev | ATATGCTAGCTTATTTGTATAACTCATCCATGCCC |
| 8296 | lrp_-36_Hind_fwd | ATATAAGCTTGTTATTGTATTTATATGAAGAAAAGGAGGCACCAC |
| 8297 | lrp-nostop_Sal_rev | ATATGTCGACTTTTGGTTGATTGTCTTCTGGTTTATCTG |
| 8435 | 020-EcoRI-N25 | GGGCCCGAATTCTCATAAAAAATTTATTTGCTTTCAGGAAAATTTTTCTG |
| 8436 | 021-GFP-stp-NheI | GGGCCCGCTAGCTTATTTGTATAGTTCATCCATGCCATGTGTAATTCC |
| 8439 | 024-SmaI-RBS-mNG | GGGCCCCCCGGGAAAAAATAAGGAGGAAAAAAAATGGTATCAAAAGGTGAAGAAGATAATATGGC |
| 8440 | 025-mNG-stp-NheI | GGGCCCGCTAGCTTATTTGTATAACTCATCCATGCCCATTAC |

| | | |
|---|---|---|
| 8441 | 026-SmaI-rbs-sfTq2ox | GGGCCCCCCGGGAAAAAATAAGGAGGAAAAAAAATGAGTAAAGGCGAAGAGTTGTTTAC |
| 8442 | 027-sfTq2ox-stp-NheI | GGGCCCGCTAGCTTATTTGTATAACTCATCCATGCCTAAAGTG |
| 8443 | 028-NheI-RBS-FP | CCCCCCGCTAGCGTTTATTAAATTAACCAACCCGGGAAAAAATAAGGAGGAAAAAAAATG |
| 8702 | P1_mNG_zapA_SalI | CTAGAGGTCGACAGGAGGACTCGGATGGTATCAAAAGG |
| 8703 | P2_mNG_zapA_EcoRI | GCGCCTGAATTCGACTTATTGCTCACGCTGCTGC |
| 8709 | P2_mNG_ftsA_KpnI | GCGCCTGGTACCGACTCATTCAAATAGAGATTTCATTAG |
| 8863 | 039-TM-NheI | GCCGGCGCTAGCTTAAGCAAACAATAAGATACCTAGTAAAAGAAC |
| 8865 | 041-tspmi-st7-sfTq2ox | GGGCCCCCCGGGAAAAATAAGGAGGAAAAAAAATGAGTAAAGGCGAAGAGTTGTTTACGGGCG |
| 8867 | 043-tspmi-st7-mNG | GGGCCCCCCGGGAAAAATAAGGAGGAAAAAAAATGGTATCAAAAGGTGAAGAAGATAATATGGCAAGC |
| 9251 | 051_ox_zip25OE-F | CTTTAGGCATGGATGAGTTATACAAAGGTGGATCAGGTGGATCAGTACCTATCCAGCGTATGAAACAGC |
| 9252 | 052_ox_zip25OE-R | GCTGTTTCATACGCTGGATAGGTACTGATCCACCTGATCCACCTTTGTATAACTCATCCATGCCTAAAG |
| 9253 | 053-Zip25-NheI | GGGCCCGCTAGCTTACTTAGGTACCCCACGTTCACCCACC |
| 9254 | 054-mNG-Zip18OE-F | GTAATGGGCATGGATGAGTTATACAAAGGTGGATCAGGTGGATCAGTACCTATCCAGCGTATGAAACAGC |
| 9255 | 055-mNG-Zip18OE-R | GCTGTTTCATACGCTGGATAGGTACTGATCCACCTGATCCACCTTTGTATAACTCATCCATGCCCATTAC |
| 9256 | 056-Zip18-NheI | GGGCCCGCTAGCTTATATCGATGAATTCCCACGTTCACC |
| 9638 | 057-tspmi-st7-FtsW | GGGCCCCCCGGGAAAAATAAGGAGGAAAAAAAATGAAGAATTTTAGAAG |
| 9639 | 058-FtsA-stp-NheI | GGGCCCGCTAGCTACGGCCGTTATCATTCAAATAGAGATTTCATTAGTTTTTTG |
| 9640 | 059-tspmi-st7-mNGecy | GGGCCCCCCGGGAAAAATAAGGAGGAAAAAAAATGGCATCATTACCAGCAACACATGAATTACATATTTTTGGTTCTATTAATGGTGTTGATTTTGATATGG |
| 9641 | 060-ZapA-stp-NheI | GGGCCCGCTAGCTACGGCCGTTATTATTGCTCACGCTGCTGCAATTTGTGAATTTGTTG |
| 9645 | 064-NGecy-lnk-PBP1-OE-F | GTAATGGGCATGGATGAGTTATACAAAGGTGGATCAGGTGGATCAGCGAAGCAAAAAATTAAAATTAAAAAAAATAAAATAG |
| 9646 | 065-NGecy-lnk-PBP1-OE-R | CTATTTTATTTTTTTTAATTTTAATTTTTTGCTTCGCTGATCCACCTGATCCACCTTTGTATAACTCATCCATGCCCATTAC |
| 9647 | 066-PBP1-stp-xbai | GGGCCCTCTAGATACGGCCGTTATTAGTCCGACTTATCCTTGTCAGTTTTAC |
| 9651 | 070-NGecy-lnk-DivIB-OE-F | GTAATGGGCATGGATGAGTTATACAAAGGTGGATCAGGTGGATCAGATGATAAAACGAAGAACGATCAACAAGAATC |
| 9652 | 071-NGecy-lnk-DivIB-OE-R | GATTCTTGTTGATCGTTCTTCGTTTTATCATCTGATCCACCTGATCCACCTTTGTATAACTCATCCATGCCCATTAC |
| 9653 | 072_DivIB-stp-NheI | GGGCCCGCTAGCTACGGCCGTTATTAATTATTCTTACTTGATTGTTTGTTAATTTTG |
| 9679 | 074-EagI-FtsW | GGGCCCCGGCCGAAAAATAAGGAGGAAAAAAAATGAAGAATTTTAGAAG |
| 9680 | 075-ox-EagI | GGGCCCCGGCCGGCTAGCTTATTTGTATAACTCATCCATGCCTAAAGTG |
| 9681 | 076-NheI-NGecy | GGGCCCGCTAGCAAAAATAAGGAGGAAAAAAAATGGCATCATTACCAGCAACACATGAATTACATATTTTTGG |
| 9729 | 077-NheI-FtsW | GGGCCCGCTAGCAAAAATAAGGAGGAAAAAAAATGAAGAATTTTAGAAG |
| 11647 | 007_st3_mNG_F | CTATCATTGATAGAGTCCCGGGTACGGCCGAAAAAAAAAATAAGGAGGAAAATGGTATCAAAAGGTGAAGAAGATAATATG |

Underlined sequences correspond to restriction sites used for cloning.

#### Supplementary References.

1. Meiresonne, N. Y. et al. Superfolder mTurquoise2 ox optimized for the bacterial periplasm allows high efficiency *in vivo* FRET of cell division antibiotic targets. Mol. Microbiol. 111, 1025–1038 (2019).

2. Goedhart, J. et al. Structure-guided evolution of cyan fluorescent proteins towards a quantum yield of 93%. Nat. Commun. 3, 751 (2012).

3. Mastop, M. et al. Characterization of a spectrally diverse set of fluorescent proteins as FRET acceptors for mTurquoise2. Sci. Rep. 7, 11999 (2017).

4. Shaner, N. C. et al. A bright monomeric green fluorescent protein derived from *Branchiostoma lanceolatum*. Nat. Methods 10, 407–9 (2013).

5. Meiresonne, N. Y., van der Ploeg, R., Hink, M. A. & den Blaauwen, T. Activity-related conformational changes in *D*,*D*-carboxypeptidases revealed by *in vivo* periplasmic förster resonance energy transfer assay in *Escherichia coli*. MBio 8, e01089-17 (2017).

6. Reichmann, N. T. et al. SEDS–bPBP pairs direct lateral and septal peptidoglycan synthesis in *Staphylococcus aureus*. Nat. Microbiol. 4, 1368–1377 (2019).

7. Meiresonne, N., Consoli, E., Mertens, L. & den Blaauwen, T. Detection of *in vivo* Protein Interactions in All Bacterial Compartments by Förster Resonance Energy Transfer with the Superfolder mTurquoise2 ox-mNeongreen FRET Pair. BIO-PROTOCOL 9, (2019).

8. Monk, I.R.; Shah, I.M.; Xu, M.; Tan, M.W.; Foster, T.J. Transforming the Untransformable: Application of Direct Transformation to Manipulate Genetically *Staphylococcus aureus* and *Staphylococcus epidermidis*. mBio, 3:e00277-11 (2012)

9. Bethesda Research Laboratories. *E. coli* DH5 alpha competent cells. Focus - Bethesda Res. Lab. 8, 9 (1986).

10. Nair, D. et al. Whole-genome sequencing of *Staphylococcus aureus* strain RN4220, a key laboratory strain used in virulence research, identifies mutations that affect not only virulence factors but also the fitness of the strain. J. Bacteriol. 193, 2332–2335 (2011).

11. Kreiswirth, B. N. et al. The toxic shock syndrome exotoxin structural gene is not detectably transmitted by a prophage. Nature 305, 709–712 (1983).

12. Fey, P. D. et al. A Genetic Resource for Rapid and Comprehensive Phenotype Screening of Nonessential *Staphylococcus aureus* Genes. MBio 4, (2013).

13. Gill, S. R. et al. Insights on evolution of virulence and resistance from the complete genome analysis of an early methicillin-resistant *Staphylococcus aureus* strain and a biofilm-producing methicillin-resistant Staphylococcus epidermidis strain. J. Bacteriol. 187, 2426–2438 (2005).

14. Pereira, P. M., Veiga, H., Jorge, A. M. & Pinho, M. G. Fluorescent reporters for studies of cellular localization of proteins in *Staphylococcus aureus*. Appl. Environ. Microbiol. 76, 4346–4353 (2010).

15. Veiga, H. et al. Cell division protein FtsK coordinates bacterial chromosome segregation and daughter cell separation in *Staphylococcus aureus*. EMBO J. 42, (2023).

16. Reed, P. et al. A CRISPRi-based genetic resource to study essential *Staphylococcus aureus* genes. MBio 15, (2024).

17. Monteiro, J. M. et al. Cell shape dynamics during the staphylococcal cell cycle. Nat. Commun. 6, 8055 (2015).

18. Pereira, A. PhD Thesis: Cell division and morphogenesis in *Staphylococcus aureus*. (2018).

19. Veiga, H., Jorge, A. M. & Pinho, M. G. Absence of nucleoid occlusion effector Noc impairs formation of orthogonal FtsZ rings during *Staphylococcus aureus* cell division. Mol. Microbiol. 80, 1366–1380 (2011).

20. Karimova, G., Pidoux, J., Ullmann, A. & Ladant, D. A bacterial two-hybrid system based on a reconstituted signal transduction pathway. Proc. Natl. Acad. Sci. U. S. A. 95, 5752–6 (1998).

21. Schäper, S. et al. Cell constriction requires processive septal peptidoglycan synthase movement independent of FtsZ treadmilling in *Staphylococcus aureus*. Nat. Microbiol. 9, 1049–1063 (2024).

22. Mertens, L. M. Y. & den Blaauwen, T. Optimising expression of the large dynamic range FRET pair mNeonGreen and superfolder mTurquoise2ox for use in the *Escherichia coli cytoplasm*. Sci. Rep. 12, 17977 (2022).
